# Bacterial anomalies in seabed sediments associated with deep water hydrocarbon seepage

**DOI:** 10.1101/2022.10.15.512386

**Authors:** Carmen Li, Oyeboade Adebayo, Deidra K. Ferguson, Scott Wang, Jayne E. Rattray, Martin Fowler, Jamie Webb, Calvin Campbell, Natasha Morrison, Adam MacDonald, Casey R.J. Hubert

## Abstract

Deep sea hydrocarbon seep detection relies predominantly on geochemical analyses of seabed marine sediment cores to identify the presence of gas or oil. The presence of seeping hydrocarbons in these locations alters resident microbial community structure, leading to culture-based biodegradation assays as a complement to geochemical tools for seep detection. Biodiversity surveys of microbial communities can offer a similar proxy for seeping hydrocarbons, but this strategy has not been extensively investigated in deep water settings. In this study, 16S rRNA gene sequencing of bacterial communities was performed on sediment cores obtained in >2500 m water depth at 43 different locations in the NW Atlantic Ocean. Core samples from as deep as 10 metres below seafloor (mbsf) were assessed for gas composition, gas isotopes and liquid hydrocarbons. Over 650 bacterial 16S rRNA gene amplicon libraries were constructed from different sediment depths at these locations. Select sites showed strong evidence for the presence of thermogenic or biogenic hydrocarbons such that bacterial population analyses revealed significant differences between hydrocarbon seep and non-seep locations. Specific bacterial indicators were associated with different sediment depth intervals. Caldatribacteriota and Campilobacterota OTUs were observed in high relative sequence abundance in hydrocarbon seep sediments, particularly in the 20-50 cmbsf interval. Furthermore, these groups were differentially abundant between sites with thermogenic and biogenic hydrocarbons. The patterns revealed here suggest that microbial screening has the potential to play a key role in hydrocarbon seep detection and characterisation in remote deep-sea environments.

## 1. Introduction

Microbiological studies of seabed hydrocarbon seeps predominantly focus on deeper anoxic sediment layers where syntrophic consortia of methanotrophic archaea and sulfate-reducing bacteria catalyse the anaerobic oxidation of methane coupled to sulfate reduction (Teske 2019). This interaction was predicted by geochemical profiles in sediments that pointed microbiologists towards these exciting discoveries of new metabolisms at sulfate-methane transition zones, often occurring a few hundred centimetres below the seafloor (cmbsf) (Beulig et al. 2019; Jørgensen et al. 2019). However, hydrocarbon seep sediments are enriched in several other taxa as well, including lineages that are poorly represented in terms of cultured isolates such as Chloroflexota, Caldatribacteriota, and candidate phyla like OD1 and OP11 (Trembath-Reichert et al. 2016; Orsi 2018; Petro et al. 2019; Dong et al. 2019; Dong et al. 2020). This reveals that there remains much more to understand about the microbial ecology in these settings.

Key processes governing distribution patterns of microorganisms in natural environments like marine sediments include environmental selection and dispersal (Hanson et al. 2012). Microbial diversity surveys that employ 16S rRNA gene amplicon sequencing tend to highlight the most abundant populations present, and thus should include key groups that actively catalyze biogeochemical processes in that environment. In this context, selection may be the strongest process affecting microbial communities in unique environments like seabed hydrocarbon seeps, differentiating these locations from the surrounding sediment. The presence of migrated hydrocarbon fluids from much deeper petroleum-bearing sediments may be expected to select for populations that directly or indirectly utilize these energy-rich compounds. In addition to exerting positive selective pressure, the input of hydrocarbons may be toxic to other marine bacteria and in parallel exert negative selection on some groups (Joye 2020). Dispersal in these kinds of deep-sea settings can be facilitated by geological upheaval forces such as those occurring at mud volcanos that reintroduce deep subsurface microbes back up to shallow sediments (Hoshino et al. 2017; Ruff et al. 2019; Rattray et al. 2022). Because of this unique combination of dispersal and selection pressures at seeps, microbiological assessments of sediments at and around these sites can potentially serve as a method for locating hydrocarbon seeps in the deep sea (Gittins et al. 2022). DNA based analyses of *in situ* microbial communities in these environments has indeed highlighted key players that are associated with the upward advection of hydrocarbon fluids (Chakraborty et al. 2020).

The aim of this study is to assess the bacterial diversity in marine sediments of the Scotian Slope in the NW Atlantic Ocean, where hydrocarbon seepage is expected to be occurring. Integrating geochemistry, microbiology and bioinformatics enabled anomalies to be identified in bacterial diversity patterns, either with distance or with depth in the seabed. Anomalies were then correlated with geochemical assessments of the presence of thermogenic or biogenic hydrocarbons in the same samples. In this study, the terms hydrocarbon-negative and hydrocarbon-positive are used in reference to attempts to identify seepage of hydrocarbons that have migrated vertically from deeper sediment intervals. In cases where geochemistry differentiates between thermogenic and biogenic hydrocarbons, this distinction is noted. Hydrocarbon negative (i.e., non-seep) sediments are ranked as such based on an absence of biogenic and thermogenic hydrocarbons that fit the above definition and may still contain hydrocarbons from recent organic matter that are mostly terrestrially derived.

## 2. Materials & Methods

### 2.1 Sample collection

Sediment coring was performed at 43 different locations along the Scotian Slope. Cores were collected aboard the CCGS Hudson in 2015 and 2016 using a piston corer with attached trigger-weight and were up to 10 meters in length (Campbell and MacDonald 2016; Campbell 2019). These were augmented by gravity cores up to 6 metres in length collected in 2018 aboard CCGS Hudson. Cores were sectioned into 1.5-meter-long intervals on board the ship and immediately inspected for signs of gas (e.g., gas cracks, bubbling, or strong odours). Sediment within 20 cm of the base of each core was immediately collected into a 500-mL IsoJar (Isotech Laboratories, Inc.), flushed with nitrogen, sealed, and stored at −20°C. The remaining full-length cores were then split longitudinally with a core splitter and further inspected for obvious hydrocarbon staining, sandy intervals, and evidence of gas. Multiple sediment samples were collected from each core for organic matter extraction, representing different intervals of interest. Samples for organic matter extraction were wrapped in aluminum foil, sealed in Whirl-pak bags, and stored at −20°C until analysis. For DNA analysis, approximately 2 mL subsamples of sediment from selected depths were collected into triplicate sterile screwcap microtubes via sterilized spatulas, and immediately frozen at −80°C on board the ship.

### 2.2 Sediment gas composition and isotope analysis

Headspace in the Isojars was analysed for gas composition as well as gas isotopes in samples with sufficient volumes of hydrocarbon gases. Aliquots were transferred to exetainers from which 0.1-1ml was removed using a Gerstel MPS2 autosampler and injected into an Agilent 7890 RGA GC (equipped with Molsieve and Poraplot Q columns). Hydrocarbons were detected using flame ionisation detection (FID), while H_2_, CO_2_, N_2_ and O_2_/Ar were measured by thermal conductivity detection (TCD).

For isotope analysis, aliquots of headspace gas were sampled with a syringe and analysed on a Trace GC2000, equipped with a Poraplot Q column, connected to a Delta plus XP isotope ratio mass spectrometer (IRMS). Repeated analyses of standards indicate that the reproducibility of δ^13^C values was better than 1 ‰ PDB (2 sigma). For some samples with low amounts of gas, preconcentration of methane was necessary prior to isotopic analysis.

Hydrogen isotopic composition of methane was determined using a GC-C-IRMS system. Headspace aliquots were sampled with a GCPal and analysed on a Trace GC2000, equipped with a Poraplot Q column, connected to a Delta plus XP IRMS. Components were decomposed to H_2_ and coke in a 1400°C furnace. The international standard NGS-2 and an in-house standard were used for testing accuracy and precision. Repeated analyses of standards indicate that the reproducibility of δD values was better than 10 ‰ PDB (2 sigma).

### 2.3 Analysis of Sediment Extractable Organic Matter

Sediment samples were analysed for total organic carbon (TOC) content, extracted and the total extract analysed by gas chromatography. Based on the amount of extract, the appearance of the gas chromatogram of the extractable organic matter (EOM), and the location of the sample, a subset (~half) of the total extracts were fractionated and analysed by GC-MS. Based on this data, hydrocarbon fractions of some samples were selected for isotopic analysis and for analysis of diamondoids.

Extraction of organic matter was performed using a Tecator Soxtec system. Crushed sediment was weighed in a pre-extracted thimble and boiled for 1 hour, then rinsed for 2 hours in 80 mL dichloromethane with 7% (vol/vol) methanol. Thimbles were pre-extracted by boiling for 10 minutes and rinsed for 20 minutes in dichloromethane with 7% (vol/vol) methanol. Copper granules, activated in concentrated hydrochloric acid, were added to react with free sulphur. To determine the amount of extractable organic matter, the weight of an aliquot of 10% of the extract was determined by transferring to a pre-weighed bottle and evaporating to dryness. For gas chromatography of the extractable organic matter, an HP7890 A instrument with a CP-Sil-5 CB-MS column (length 30 m, i.d. 0.25 mm, film thickness 0.25 μm) was used and C_20_D_42_ was used as an internal standard. The temperature programme was 50°C (1 min) - 4°C/min - 320°C (25 min).

Samples selected for more detailed analysis were fractionated using a MPLC system constructed similar to that described by Radke (1980). The system included two HPLC pumps, sample injector, sample collector and two packed columns. The precolumn was filled with Kieselgel 100, heated at 600°C for 2 hours to deactivate the silica. The main column, a LiChroprep Si60 column, was heated at 120°C for 2 hours with a helium flow to remove water. Approximately 30 mg of deasphaltened oil or EOM diluted in 1 ml hexane was injected into a sample loop. The solvents used were hexane and dichloromethane with saturated hydrocarbons, aromatic hydrocarbons and polars (NSO-fraction) collected. The hydrocarbon fractions were analyzed for GC-MS using a Thermo Scientific DFS high resolution instrument. This was tuned to a resolution of 3000 with a 60 m CP-Sil-5 CB-MS column (length 50 m, i.d. 0.25 mm, film thickness 0.25 μm) used and data was acquired in Selected Ion Recording (SIR) mode. The temperature programme was 50°C (1 min) - 20°C/min - 120°C - 2°C/min - 320 °C (20 min). D_4_-27ααR was used as an internal standard for saturated compounds; D_8_-Naphthalene, D_10_-Biphenyl, D_10_-Phenanthrene and D_12_-Chrysene were used as internal standards for aromatic compounds.

### 2.4 GCMS of diamondoids in saturated fractions

A Thermo Scientific TSQ Quantum XLS instrument tuned to a resolution of 0.4 mass units, with a CP-Sil-5 CB-MS column (60 m length, i.d. 0.25 mm, film thickness 0.25μm) was used and data was acquired in Selected Ion Recording (SIR) mode. The temperature programme was 50 °C (1 min) - 3 °C/min. - 230 °C - 15 °C/min - 325 °C (20 min). D16-Adamantane and D3-1-Methyl-Diamantane were used as internal standards.

### 2.5 DNA analysis

Genomic DNA was extracted from ~500 mg of individual triplicate sediment subsamples using the DNEasy PowerSoil kit (Qiagen), following a slightly modified version of the manufacturer’s protocol. Bacterial 16S rRNA genes were amplified from the resulting genomic DNA using 1-10 ng genomic DNA, KAPA HiFi HotStart Ready mastermix (Roche), and 0.1 uM (final concentration) of bacteria 16S rRNA gene-specific primers modified with Illumina adapters (S-D-Bact-0341-a-S-17 5’-TCGTCGGCAGCGTCAGATGTGTATAAGAGACAGC<u>CTACGGGNGGCWGCAG</u>-3’and S-D-Bact-0785-a-A-21 5’-GTCTCGTGGGCTCGGAGATGTGTATAAGAGACAG<u>GACTACHVGGGTATCTAATCC</u>-3’). Each of the triplicate DNA extractions was amplified in triplicate PCR reactions using a program with an initial denaturation at 95°C for 5 minutes, followed by 10 cycles (with a decreasing annealing temperature by 1°C per cycle) of 95°C for 30 seconds, 60 - 51°C for 45 seconds, and 72°C for 1 minute, then 20 cycles of 95°C for 30 seconds, 55°C for 45 seconds, and 72°C for 1 minute, and a final extension at 72°C for 5 minutes. PCR reactions were then prepared into amplicon libraries for sequencing following an adapted version of the Illumina 16S Metagenomic Sequencing Library Preparation protocol (Illumina Inc.). Libraries were quantified using the Qubit dsDNA HS Assay kit on a Qubit 2.0 fluorometer (ThermoFisher Scientific), normalized, pooled, and verified on a 2100 Bioanalyzer System (Agilent Technologies, Inc.) before being sequenced on an inhouse MiSeq benchtop sequencer using a 600-cycle MiSeq Reagent V3 kit (Illumina, Inc.). Raw sequence data was deposited in the NCBI BioProject database with accession code PRJNA889844 under accession numbers SAMN31261098 to SAMN31261765.

In total 668 amplicon libraries were sequenced (i.e., approximately 224 different samples in triplicate). OTU analysis at a 97% sequence similarity cut-off was performed using MetaAmp (Dong et al. 2017). All sequences were trimmed to a fixed length of 350 bases and identity assigned based on SILVA (version 138). Non-metric multidimensional scaling (NMDS) based on Bray-Curtis dissimilarity was conducted using MetaAmp, and IndicSpecies (Cáceres and Legendre 2009) and ANOSIM (Oksanen et al., 2020) analyses were conducted in R (Version 4.0.2).

IndicSpecies analysis was performed with samples grouped into hydrocarbon-positive (seep) and hydrocarbon-negative (non-seep) categories, as defined by the presence/absence of geochemical indicators described above for each of the 43 sediment cores. Additional IndicSpecies analysis further categorized hydrocarbon-positive samples as being either thermogenic or biogenic (the latter being due to the presence of methane derived from microbial methanogenesis in relatively shallow sediments), enabling comparison of thermogenic and biogenic sites with each other and with hydrocarbon-negative samples (i.e., non-seep sites lacking thermogenic and biogenic hydrocarbon advection). Samples from sites with inconclusive geochemical evidence for thermogenic or biogenic hydrocarbons were removed prior to IndicSpecies analyses. Only those OTUs present in at least 5% relative sequence abundance in at least one sample were included in these analyses. IndicSpecies analysis was performed on samples from all depths and on samples grouped into depth interval categories of 0 cmbsf, 20-50 cmbsf, and >50 cmbsf. Resulting indicator OTUs, based on a significance level of 0.05, were further investigated if they showed a significant increase (based on a two-sample T-test assuming unequal variance) of >5% in average relative sequence abundances when comparing any of the hydrocarbon-positive categories (thermogenic, biogenic or combined thermogenic and biogenic) with hydrocarbon-negative samples, in at least one of the depth interval categories 0 cmbsf, 20-50 cmbsf and >50 cmbsf.

ANOSIM was performed for pairwise comparisons of samples grouped into hydrocarbon-negative, hydrocarbon-positive (thermogenic or biogenic), thermogenic and biogenic categories. For each ANOSIM, all sediment depths were included in a single analysis of negative or positive samples. Subsequently, samples from each of 0 cmbsf, 20-50 cmbsf, or >50 cmbsf intervals were analyzed in negative and positive categories. Samples from sites with inconclusive geochemical evidence for thermogenic hydrocarbons were compared individually against hydrocarbon-negative, thermogenic and biogenic categories, with samples of all depths as a single analysis as well as samples of individual depths as separate analyses.

## 3. Results

### 3.1 Geochemical analyses identify hydrocarbon seep and non-seep locations

Sediments from the bottom of each piston core were used for site ranking and were assessed for biogenic and/or thermogenic hydrocarbons based on analysis of gas composition and isotopes, saturate and aromatic hydrocarbons within extracted organic matter and their carbon isotopes, and diamondoids (Figure 1). Out of the 43 sampling locations, four sites showed strong signals for seepage of either biogenic gas or a mix of biogenic and thermogenic gaseous and liquid hydrocarbons (Table 1).

**Figure 1.**
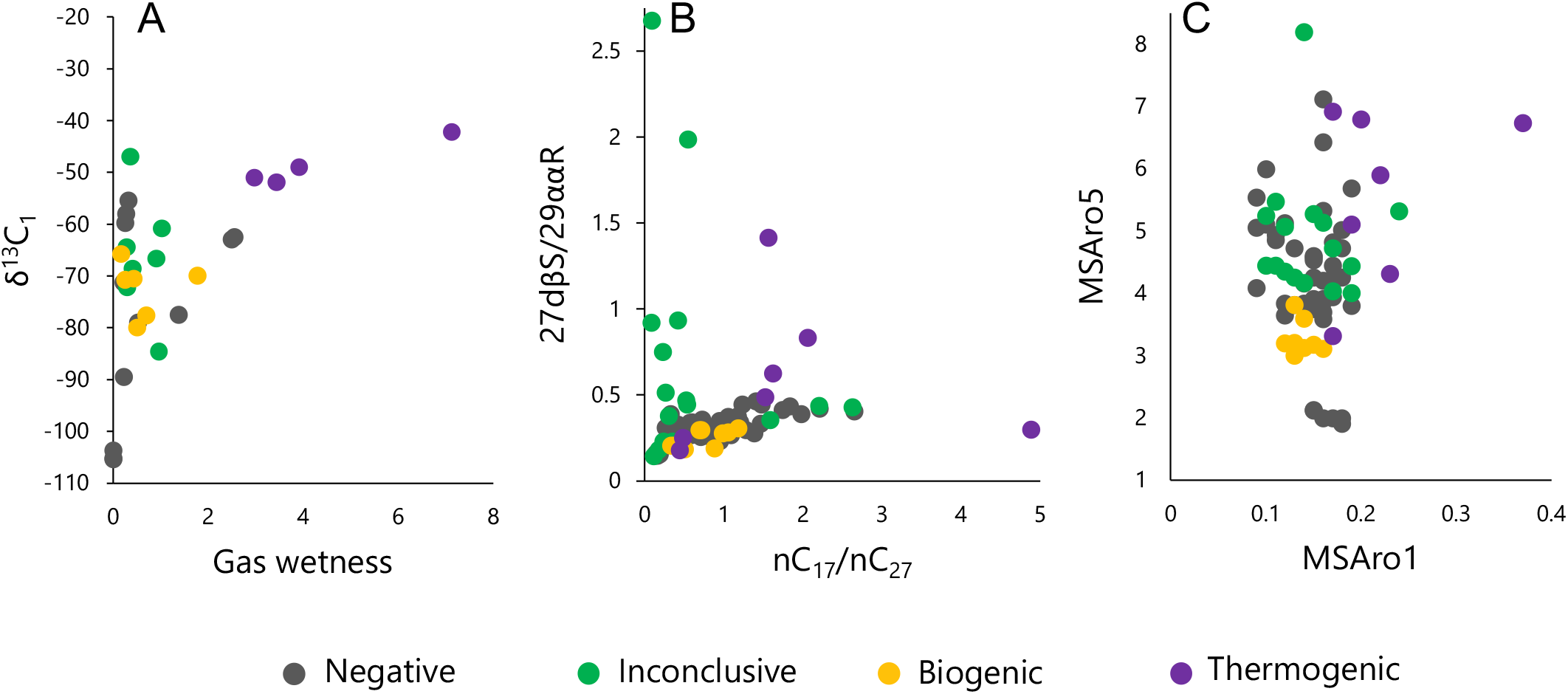
Select hydrocarbon geochemistry parameters measured on Scotian Slope sediment cores: **A.** δ^13^C-CH_4_ versus gas wetness (n=31 cores), **B.** nC_17_/nC_27_ versus 27dβS/29ααR (n=75 sediment samples from 42 cores), and **C.** MSAro1 versus MSAro5 (n=75 sediment samples from 42 cores).

**Table 1.** Select geochemistry parameters measured on Scotian Slope sediments. Values obtained for gas isotopes (δ^13^C-CH4), gas wetness, alkanes (n-C_17_/nC_17_+nC_27_), and steranes (27dβS/ C29ααR^1^, MSAro1^2^, and MSAro5^3^) are listed for 43 sites. Empty cells signify that samples were not analyzed for that respective parameter.

| Site | Upper Depth (m) | Lower Depth (m) | $\delta^{13}\text{C-CH}_4$ | Gas Wetness ($\text{C}_2\text{-C}_4/\text{C}_1\text{-C}_4$) | $\text{n-C}_{17}/(\text{n-C}_{17}+\text{n-C}_{27})$ | $27\text{d}\beta\text{S}/\text{C}_{29\alpha\alpha}\text{R}$ | MSAro1 | MSAro5 | Hydrocarbon Ranking |
| --- | --- | --- | --- | --- | --- | --- | --- | --- | --- |
| 15-3 | 1.9 | 1.96 |  |  | 0.12 |  |  |  | Negative |
|  | 2.48 | 2.54 |  |  | 0.15 |  |  |  |  |
|  | 4.85 | 4.91 |  |  | 0.15 |  |  |  |  |
|  | 5.11 | 5.16 |  | 0 |  |  |  |  |  |
| 15-5 | 2.59 | 2.64 |  |  | 0.17 |  |  |  | Negative |
|  | 4.14 | 4.2 |  |  | 0.13 |  |  |  |  |
|  | 5.74 | 5.81 |  |  | 0.2 |  |  |  |  |
|  | 6.22 | 6.31 |  |  | 0.21 | 0.31 | 0.15 | 4.53 |  |
|  | 7.39 | 7.44 | -89.5 | 0.16 |  |  |  |  |  |
|  | 7.44 | 7.49 |  | 0.22 |  |  |  |  |  |
| 15-6 | 2.2 | 2.26 |  |  | 0.1 |  |  |  | Inconclusive |
|  | 2.98 | 3.04 | -84.6 | 0.22 |  |  |  |  |  |
|  | 3.28 | 3.33 |  |  | 0.26 |  |  |  |  |
|  | 3.6 | 3.65 |  |  | 0.15 |  |  |  |  |
|  | 5 | 5.05 |  |  | 0.19 |  |  |  |  |
|  | 5.61 | 5.66 | -66.6 | 0.96 |  |  |  |  |  |
|  | 5.66 | 5.71 |  | 0.91 |  |  |  |  |  |
|  | 2.2 | 2.26 |  |  | 0.10 | 0.14 | 0.12 | 4.34 |  |
|  | 3.28 | 3.33 |  |  | 0.26 | 0.23 | 0.13 | 4.25 |  |
|  | 3.6 | 3.65 |  |  | 0.15 | 0.18 | 0.19 | 4.00 |  |
|  | 5 | 5.05 |  |  | 0.19 | 0.23 | 0.17 | 4.03 |  |
| 15-8 | 2.96 | 3.01 |  |  | 0.17 |  |  |  | Negative |
|  | 5.75 | 5.8 |  |  | 0.12 |  |  |  |  |
|  | 7.39 | 7.44 |  |  | 0.3 | 0.33 | 0.14 | 4.15 |  |
|  | 10.05 | 10.1 |  | 0 |  |  |  |  |  |
| 15-9 | 1.44 | 1.49 |  |  | 0.19 | 0.75 | 0.11 | 5.47 | Inconclusive |
|  | 2 | 2.06 |  |  | 0.08 | 2.68 | 0.11 | 4.44 |  |
|  | 2.97 | 3.02 |  |  | 0.34 | 0.47 | 0.16 | 5.13 |  |
|  | 3.63 | 3.69 |  |  | 0.23 | 0.38 | 0.14 | 4.16 |  |
|  | 4.28 | 4.33 | -60.8 | 1.03 |  |  |  |  |  |
|  | 4.33 | 4.38 | -68.6 | 0.41 |  |  |  |  |  |
| 15-10 | 3.72 | 3.77 |  |  | 0.09 |  |  |  | Negative |
|  | 4.6 | 4.65 |  |  | 0.36 | 0.34 | 0.14 | 3.83 |  |
|  | 4.81 | 4.86 |  | 0 |  |  |  |  |  |
| 15-14 | 1.8 | 1.85 |  |  | 0.13 |  |  |  | Negative |
|  | 4.3 | 4.35 |  |  | 0.39 |  |  |  |  |
|  | 6.55 | 6.6 |  |  | 0.48 | 0.32 | 0.16 | 4.20 |  |
|  | 7.79 | 7.94 |  |  | 0.55 | 0.44 | 0.15 | 4.59 |  |
| 15-15 | 2.8 | 2.85 |  |  | 0.12 |  |  |  | Negative |
|  | 4.3 | 4.35 |  |  | 0.22 |  |  |  |  |
|  | 5.3 | 5.35 |  |  | 0.26 |  |  |  |  |
| 15-18 | 2.65 | 2.7 |  |  | 0.12 |  |  |  | Negative |
|  | 3.04 | 3.09 |  |  | 0.12 | 0.16 | 0.17 | 3.93 |  |
|  | 3.39 | 3.54 |  |  | 0.18 |  |  |  |  |
|  | 3.68 | 3.73 |  |  | 0.12 |  |  |  |  |
|  | 3.89 | 3.98 |  | 0 |  |  |  |  |  |
|  | 4.23 | 4.31 |  |  | 0.17 |  |  |  |  |
|  | 4.66 | 4.73 |  |  | 0.19 |  |  |  |  |
|  | 5.41 | 5.46 |  |  | 0.18 |  |  |  |  |
|  | 7.68 | 7.73 |  | 0 |  |  |  |  |  |
| 15-19 | 2 | 2.05 |  |  | 0.32 |  |  |  | Negative |
|  | 4.56 | 4.61 |  |  | 0.12 |  |  |  |  |
|  | 5.23 | 5.28 |  |  | 0.13 |  |  |  |  |
| 15-21 | 1.54 | 1.59 |  |  | 0.3 |  |  |  | Negative |
|  | 2.95 | 3 |  |  | 0.21 |  |  |  |  |
|  | 4.47 | 4.53 |  |  | 0.35 |  |  |  |  |
|  | 6.81 | 6.86 |  | 0 |  |  |  |  |  |
| 16-4 | 1.31 | 1.36 |  |  | 0.1 |  |  |  | Inconclusive<br>-<br>thermogenic |
|  | 1.61 | 1.66 | -47 |  |  |  |  |  |  |
| 16-5 | 1.88 | 1.93 |  |  | 0.16 |  |  |  | Negative |
|  | 3.6 | 3.65 |  |  | 0.16 |  |  |  |  |
|  | 4.35 | 4.42 |  |  | 0.19 |  |  |  |  |
|  | 4.86 | 4.91 |  |  | 0.16 |  |  |  |  |
|  | 6.09 | 6.14 |  |  | 0.25 | 0.39 | 0.12 | 3.83 |  |
| 16-6 | 0.15 | 0.2 |  |  | 0.14 |  |  |  | Negative |
|  | 2.24 | 2.29 |  |  | 0.08 |  |  |  |  |
|  | 5.71 | 5.76 |  |  | 0.17 |  |  |  |  |
|  | 6.02 | 6.07 |  |  | 0.16 | 0.17 | 0.09 | 4.08 |  |
|  | 6.07 | 6.12 |  |  | 0.19 |  |  |  |  |
|  | 8.02 | 8.07 | -59.8 |  |  |  |  |  |  |
| 16-8 | 2.8 | 2.85 |  |  | 0.14 |  |  |  | Negative |
|  | 5.78 | 5.83 |  |  | 0.24 | 0.30 | 0.11 | 4.93 |  |
| 16-13 | 1.81 | 1.86 |  |  | 0.25 |  |  |  | Negative |
|  | 2.55 | 2.6 |  |  | 0.19 |  |  |  |  |
|  | 3.4 | 3.45 |  |  | 0.22 |  |  |  |  |
| 16-14 | 2.95 | 3 |  |  | 0.16 |  |  |  | Negative |
|  | 3.75 | 3.85 |  |  | 0.16 |  |  |  |  |
|  | 3.99 | 4.04 |  |  | 0.16 |  |  |  |  |
|  | 5.73 | 5.78 |  |  | 0.17 |  |  |  |  |
|  | 7.75 | 7.8 |  |  | 0.19 |  |  |  |  |
|  | 8.71 | 8.76 |  |  | 0.17 |  |  |  |  |
|  | 9.33 | 9.38 | -58 |  |  |  |  |  |  |
| 16-15 | 6.09 | 6.14 |  |  | 0.16 |  |  |  | Negative |
|  | 6.58 | 6.63 |  |  | 0.18 |  |  |  |  |
|  | 6.84 | 6.94 |  |  | 0.26 |  |  |  |  |
|  | 8.9 | 8.95 |  |  | 0.19 |  |  |  |  |
| 16-16 | 0.98 | 1.03 |  |  | 0.23 |  |  |  | Negative |
|  | 1.51 | 1.56 |  |  | 0.26 |  |  |  |  |
|  | 2.92 | 2.97 |  |  | 0.15 |  |  |  |  |
| 16-18 | 0.87 | 0.92 |  |  | 0.29 |  |  |  | Negative |
|  | 1.29 | 1.34 |  |  | 0.27 |  |  |  |  |
|  | 1.41 | 1.46 |  |  | 0.2 |  |  |  |  |
|  | 1.58 | 1.63 |  |  | 0.21 |  |  |  |  |
|  | 2 | 2.09 | -77.5 |  |  |  |  |  |  |
|  | 2.9 | 2.95 | -79 |  |  |  |  |  |  |
| 16-19 | 1.43 | 1.48 |  |  | 0.23 |  |  |  | Negative |
|  | 2.36 | 2.41 |  |  | 0.31 |  |  |  |  |
|  | 3.35 | 3.4 |  |  | 0.41 |  |  |  |  |
| 16-21 | 0.12 | 0.17 |  |  | 0.27 |  |  |  | Inconclusive<br>- biogenic<br>and<br>thermogenic |
|  | 1.48 | 1.55 |  |  | 0.21 | 0.51 | 0.12 | 5.06 |  |
|  | 1.49 | 1.58 |  |  | 0.30 | 0.93 | 0.10 | 5.24 |  |
|  | 2.1 | 2.15 |  |  | 0.27 |  |  |  |  |
|  | 2.3 | 2.35 |  |  | 0.35 | 1.98 | 0.24 | 5.31 |  |
|  | 2.36 | 2.41 | -64.4 |  |  |  |  |  |  |
|  | 2.5 | 2.55 |  |  | 0.35 | 0.44 | 0.14 | 8.19 |  |
|  | 2.85 | 2.9 |  |  | 0.33 |  |  |  |  |
|  | 3.31 | 3.36 | -72.2 |  |  |  |  |  |  |
| 16-24 | 5.1 | 5.15 |  |  | 0.59 | 0.33 | 0.11 | 4.85 | Negative |
|  | 6.95 | 7 |  |  | 0.44 |  |  |  |  |
| 16-26 | 1.89 | 1.94 |  |  | 0.46 | 0.25 | 0.12 | 3.64 | Negative |
|  | 4.83 | 4.88 |  |  | 0.3 |  |  |  |  |
|  | 7.79 | 7.84 |  |  | 0.73 | 0.40 | 0.17 | 4.24 |  |
| 16-29 | 4.85 | 4.9 |  |  | 0.43 |  |  |  | Negative |
|  | 6.42 | 6.48 |  |  | 0.22 |  |  |  |  |
|  | 7.04 | 7.09 |  |  | 0.30 | 0.25 | 0.18 | 4.72 |  |
| 16-30 | 3.29 | 3.34 |  |  | 0.47 |  |  |  | Negative |
|  | 7.88 | 7.93 |  |  | 0.60 | 0.44 | 0.16 | 6.42 |  |
| 16-32 | 2.6 | 2.65 |  |  | 0.72 | 0.43 | 0.19 | 4.43 | Inconclusive<br>-<br>thermogenic |
|  | 3.85 | 3.9 |  |  | 0.61 | 0.35 | 0.15 | 5.27 |  |
|  | 5.4 | 5.45 |  |  | 0.69 | 0.43 | 0.17 | 4.72 |  |
| 16-33 | 6.1 | 6.15 |  |  | 0.39 |  |  |  | Negative |
| 16-35 | 1.5 | 1.55 |  |  | 0.58 | 0.46 | 0.16 | 5.32 | Negative |
| 16-37 | 2.7 | 2.75 |  |  | 0.26 |  |  |  | Negative |
| 16-38 | 5.63 | 5.68 |  |  | 0.41 | 0.25 | 0.13 | 3.81 | Negative |
| 16-40 | 5.55 | 5.6 |  |  | 0.24 |  |  |  | Negative |
| 16-41 | 0.53 | 0.6 |  |  | 0.33 | 0.25 | 0.17 | 6.91 | Positive<br>-<br>thermogenic |
|  | 2.08 | 2.13 |  |  | 0.31 | 0.18 | 0.20 | 6.79 |  |
|  | 2.27 | 2.32 |  |  | 0.61 | 1.41 | 0.37 | 6.73 |  |
|  | 3.1 | 3.15 |  |  | 0.83 | 0.30 | 0.22 | 5.89 |  |
|  | 3.32 | 3.37 | -42.2 |  |  |  |  |  |  |
|  | 3.37 | 3.44 | -49 |  |  |  |  |  |  |
| 16-42 | 7.18 | 7.23 |  |  | 0.3 |  |  |  | Negative |
| 16-44 | 7.55 | 7.6 |  |  | 0.29 |  |  |  | Negative |
| 16-45 | 4.9 | 4.95 |  |  | 0.41 |  |  |  | Negative |
| 16-48 | 0.8 | 0.86 |  |  | 0.36 |  |  |  | Positive<br>- biogenic |
|  | 1.12 | 1.17 |  |  | 0.33 |  |  |  |  |
|  | 1.48 | 1.55 |  |  | 0.42 |  |  |  |  |
|  | 1.64 | 1.7 |  |  | 0.33 | 0.18 | 0.13 | 2.99 |  |
|  | 2.32 | 2.39 |  |  | 0.47 |  |  |  |  |
|  | 5.01 | 5.06 | -77.7 |  |  |  |  |  |  |
|  | 5.28 | 5.33 | -70.8 |  |  |  |  |  |  |
|  | 5.4 | 5.45 |  |  | 0.27 |  |  |  |  |
|  | 5.75 | 5.8 |  |  | 0.42 | 0.29 | 0.15 | 3.17 |  |
|  | 7.11 | 7.16 |  |  | 0.51 | 0.28 | 0.13 | 3.20 |  |
|  | 8.51 | 8.56 | -70.5 |  |  |  |  |  |  |
| 16-49 | 0.77 | 0.82 |  |  | 0.33 |  |  |  | Positive<br>- biogenic |
|  | 1.44 | 1.49 |  |  | 0.54 | 0.31 | 0.16 | 3.10 |  |
|  | 2.26 | 2.31 |  |  | 0.47 | 0.19 | 0.12 | 3.19 |  |
|  | 2.87 | 2.92 | -80 |  |  |  |  |  |  |
|  | 3.62 | 3.67 |  |  | 0.2 |  |  |  |  |
|  | 4.03 | 4.09 |  |  | 0.41 | 0.29 | 0.13 | 3.81 |  |
|  | 4.43 | 4.48 |  |  | 0.25 | 0.20 | 0.14 | 3.12 |  |
|  | 4.49 | 4.54 | -65.8 |  |  |  |  |  |  |
|  | 4.6 | 4.65 |  |  | 0.27 | 0.20 | 0.13 | 3.18 |  |
|  | 6.2 | 6.25 |  |  | 0.50 | 0.27 | 0.14 | 3.59 |  |
|  | 6.4 | 6.45 | -70 |  |  |  |  |  |  |
| 18-6 | 1.32 | 1.37 |  |  | 0.29 |  |  |  | Negative |
|  | 2.86 | 2.91 |  |  | 0.37 |  |  |  |  |
|  | 3.83 | 3.88 |  |  | 0.28 |  |  |  |  |
|  | 4.51 | 4.56 |  | 0 |  |  |  |  |  |
| 18-7 | 0.38 | 0.43 |  | 2.97 | 0.6 | 0.18 | 0.17 | 3.31 | Positive<br>-<br>thermogenic |
|  | 0.34 | 0.38 |  | 3.44 | 0.62 | 0.40 | 0.23 | 4.31 |  |
|  | 0.32 | 0.34 |  |  | 0.67 | 0.35 | 0.19 | 5.10 |  |
| 18-14 | 4.19 | 4.27 |  |  | 0.21 |  |  |  | Negative |
|  | 4.33 | 4.38 |  |  | 0.18 |  |  |  |  |
|  | 4.88 | 4.93 |  |  | 0.22 | 0.09 | 0.15 | 2.12 |  |
|  | 5.66 | 5.71 |  | 0 |  |  |  |  |  |
| 18-15 | 2.85 | 2.9 |  |  | 0.15 |  |  |  | Negative |
|  | 3.28 | 3.33 |  |  | 0.21 |  |  |  |  |
|  | 4.66 | 4.71 |  |  | 0.49 | 0.13 | 0.16 | 1.99 |  |
|  | 5.91 | 5.96 |  | 0 |  |  |  |  |  |
| 18-22 | 2.73 | 2.78 |  |  | 0.35 |  |  |  | Negative |
|  | 3.65 | 3.7 |  |  | 0.32 |  |  |  |  |
|  | 4.14 | 4.19 |  |  | 0.37 |  |  |  |  |

<sup>1</sup>A ratio of C<sub>27</sub> 13β(H),17α(H) 20S diasterane/ C<sub>29</sub> 5α(H),14α(H),17α(H) 20R sterane measured using m/z 217 mass chromatogram
<sup>2</sup>(C<sub>21</sub>MA+C<sub>22</sub>MA)/(C<sub>21</sub>MA+C<sub>22</sub>MA+βSC<sub>27</sub>MA+βRC<sub>27</sub>MA+βRC<sub>27</sub>DMA+αSC<sub>27</sub>MA+βSC<sub>28</sub>MA+βSC<sub>28</sub>DMA+αRC<sub>27</sub>DMA+αSC<sub>27</sub>DMA+αRC<sub>27</sub>MA+αSC<sub>28</sub>MA+αSC<sub>29</sub>MA+αRC<sub>29</sub>MA) measured using m/z 253 mass chromatogram
<sup>3</sup>(2,6-Dimethylnaphthalene + 2,7- Dimethylnaphthalene)/1,5- Dimethylnaphthalene measured using m/z 156 mass chromatogram

Samples from sites 16-41 and 18-7 contained strong indicators of thermogenic hydrocarbons, including a thermogenic gas hydrate, H_2_S gas and C_1_-C_4_ hydrocarbon gases with δ^13^C values indicative of an oil-associated thermogenic origin (Fig. 1A). Samples from both sites also showed higher amounts of C_15_-C_20_ n-alkanes relative to plant-derived C_21_-C_33_ odd-numbered n-alkanes compared to other samples (Table 1), and higher abundance of short-chain steranes and diasteranes (Table 1) indicating more mature thermogenic contribution. Similarly, diamondoids were detected in high concentrations at these two sites (Table 1).

The highest total hydrocarbon gases (THCG) were measured from sites 16-48 and 16-49 where gas hydrates consisting of predominantly methane (99.6-99.7%) were recovered. This methane was isotopically light (−71.93 ‰ to −73 ‰), indicative of a biogenic origin (Table 1). There was no indication from geochemical analyses for thermogenic liquid hydrocarbons in samples from these cores. Therefore hydrocarbons at these sites are predominantly derived from microbial methanogenesis.

### 3.2 Geochemical evidence is inconclusive for thermogenic hydrocarbon seepage in some locations

Evidence for the presence of thermogenic hydrocarbons was inconclusive in five of the 43 locations sampled (Table 1). In these cases, some parameters suggested the absence of thermogenic or biogenic hydrocarbon seepage, while other parameters suggested the presence of thermogenic hydrocarbons, as outlined below. In these instances, based on geochemistry alone, the possibility of thermogenic and/or biogenic hydrocarbons cannot not be ruled out.

Inconclusive sites 15-6 and 15-9 had C_2_-C_4_ gases with isotopic signatures suggestive of mixed biogenic-thermogenic origin. Organic matter extracts from from 15-9 samples showed a secondary unresolved complex mixture (UCM) under nC_23_-nC_27_ alkanes and a higher abundance of diasteranes, providing possible evidence of thermogenic hydrocarbons. A predominance of C_27_ over C_29_ steranes in these sediments, as well as higher abundances of tricyclic terpanes relative to hopane and of rearranged steranes and hopanes, is similar to thermogenic site 16-41 referred to above. Sediment from site 15-9 additionally contained diamondoids, which indicated a thermal maturity of the organic matter much higher than suggested by other compounds.

Inconclusive site 16-21 had similarly conflicting results, including methane that was isotopically light (i.e. outside the thermogenic range; Bernard et al. 1978; Whiticar. 1994) and low abundances of lighter hydrocarbons (<nC_20_) relative to the higher land plant-derived C_25_-C_31_ n-alkanes. On the other hand, 16-21 samples have higher concentrations of short-chain steranes and diasteranes, and a predominance of C_27_ over C_29_ diasteranes. Gas samples from this site also contained H_2_S. The presence of liquid thermogenic hydrocarbons at 16-21 thus cannot be ruled out.

Dry gas with isotopically heavy methane was recovered from site 16-4 suggestive of thermogenic gas seepage. Samples from 16-32 had a high proportion of C_15_-C_20_ n-alkanes relative to plant-derived odd-numbered C_23_-C_31_ n-alkanes. Extracted organic matter from 16-32 consisted mostly of hydrocarbons with a high saturate/aromatic ratio. EOM-GCs of these samples suggest a component of higher maturity hydrocarbons. Relatively high levels of diamondoids were also measured at 16-32.

### 3.3 Bacterial community composition varies with sediment depth and presence of hydrocarbon seepage

Bacterial diversity profiles based on 16S rRNA gene amplicon libraries clustered consistently when grouped by sediment depth intervals of 0 cmbsf, 20-50 cmbsf, and >50 cmbsf (Figure 2). This suggests that surface sediment communities are distinct from subsurface communities, independent of site location and the presence of hydrocarbon seepage.

**Figure 2.**
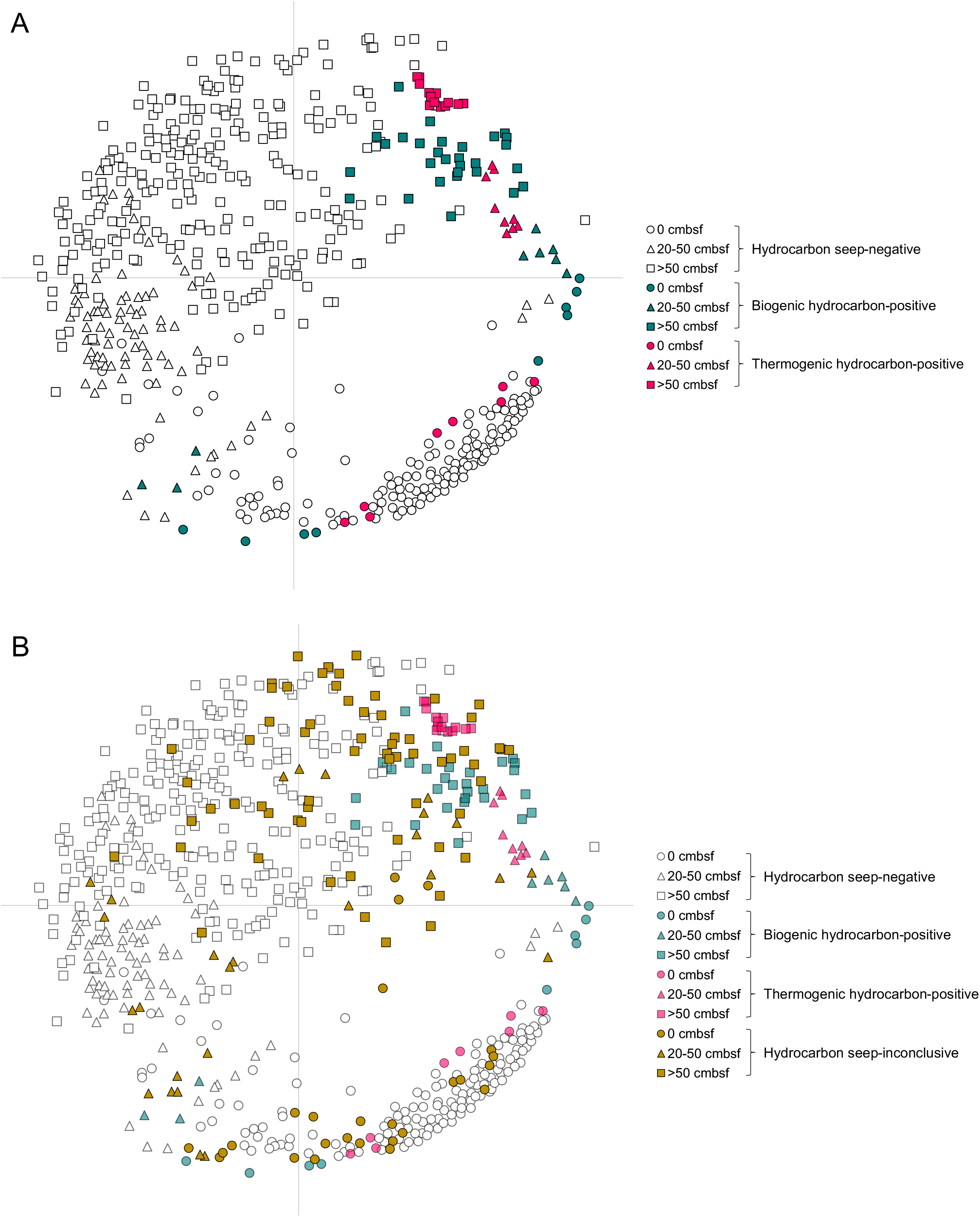
Non-metric multidimensional scaling of Bray-Curtis dissimilarity among 668 bacterial communities in samples from different sediment depths from 43 Scotian Slope sediment cores. The hydrocarbon ranking at a sediment core location is indicated by different colours for the presence of thermogenic hydrocarbon seepage (magenta), biogenic hydrocarbon seepage (teal), the absence of biogenic or thermogenic hydrocarbon seepage (white) or cores for which the presence of thermogenic or biogenic hydrocarbon seepage was inconclusive (gold). The shape of the symbols indicates the sample depth as being from 0 cmbsf (circles), 20-50 cmbsf (triangles), or >50 cmbsf (squares). **A**. Samples with inconclusive evidence for hydrocarbon seepage are omitted on the plot for easier visual comparison of hydrocarbon seepage-positive and hydrocarbon seepage-negative samples (n=549). **B.** All samples are included in the plot (n=668).

Within this overall trend, some clear outliers were observed. These were typically from the 20-50 cmbsf depth interval at sites with thermogenic or biogenic hydrocarbons, which clustered with >50 cmbsf samples from these same hydrocarbon seep locations (Figure 2). The >50 cmbsf samples from hydrocarbon seep sites clustered tightly within a more disparate group of >50 cmbsf non-seep samples, suggesting that bacterial communities deeper in the subsurface where there is hydrocarbon seepage may have unique features while still exhibiting an overall signature of deeper subsurface bacterial communities. This pattern suggests that the bacterial communities in seep sediments differ most noticeably from non-seep sediments in the 20-50 cmbsf range. Despite this observation, ANOSIM comparisons indicate that the differences between seep and non-seep sediments are significant for all three depth intervals (0 cmbsf, 20-50 cmbsf, and >50 cmbsf) (Table S8).

Hydrocarbon seep sediments further separate into thermogenic and biogenic groups at the 20-50 cmbsf and >50 cmbsf depth intervals. This suggests that bacterial community compositions are distinct from one another in sediments at these depths based on whether the presence of hydrocarbons is derived from a biogenic or thermogenic origin.

### 3.4 Caldatribacteriota and Campilobacterota OTUs in different sediment depth intervals are indicative of migrated thermogenic hydrocarbons

IndicSpecies analysis highlighted 5 OTUs with elevated relative sequence abundance in thermogenic hydrocarbon seep sediments at different depth intervals (Figure 3). Notably, Caldatribacteriota OTUs 1, 3029, 21130, Campilobacterota OTU 8, and Acidobacteriota OTU 21 were found in significantly higher average relative sequence abundances (>5% in at least one depth category) in cores with thermogenic hydrocarbons compared to non-seep sediment cores (Table S1). These five OTUs alone constitute 5% - 57% of the overall bacterial communities when thermogenic hydrocarbons are present (Figure 3).

**Figure 3.**
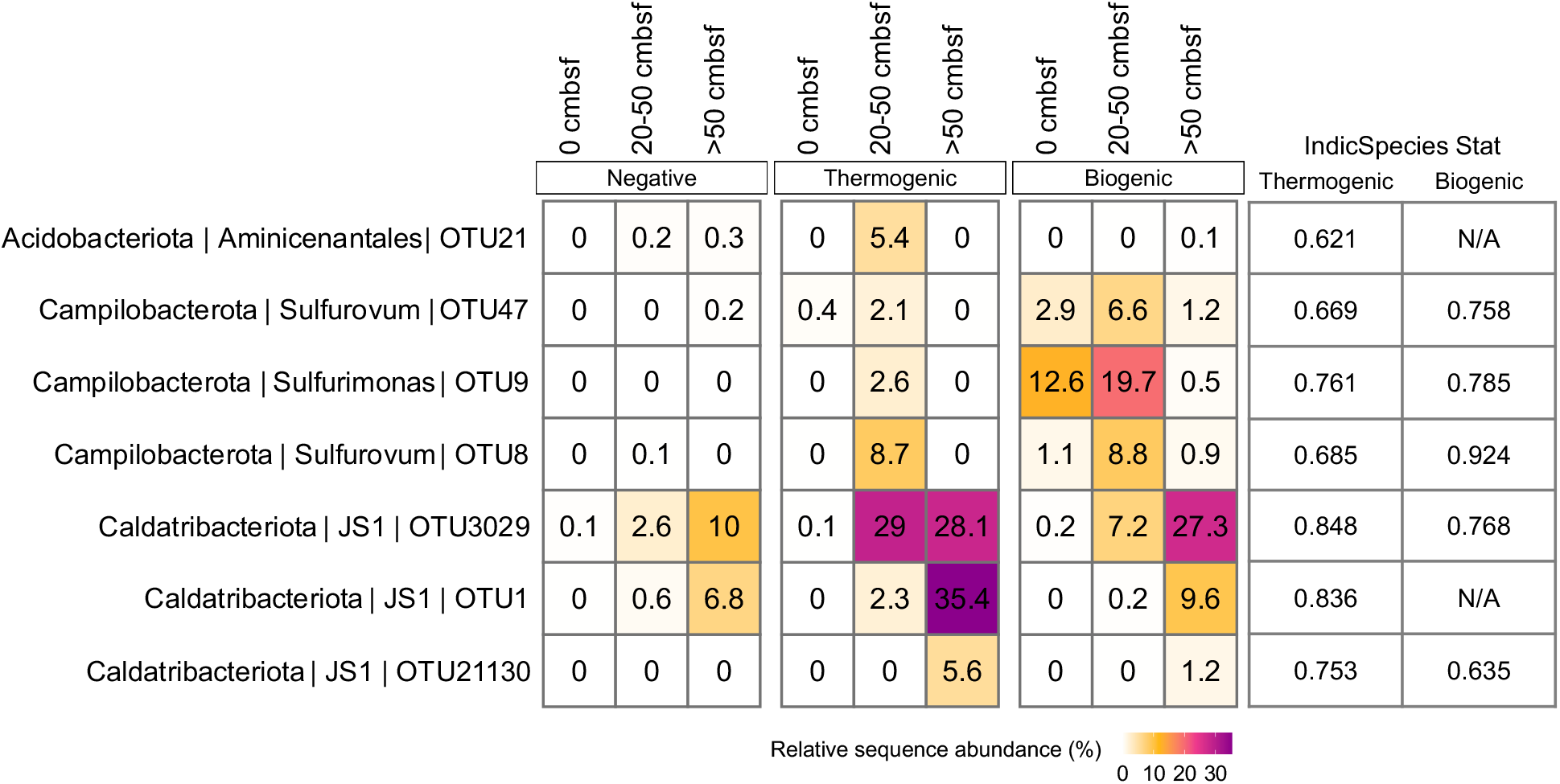
Average relative sequence abundances of OTUs identified as indicators for thermogenic and biogenic hydrocarbon seepage by IndicSpecies analysis in non-seep, thermogenic seep and biogenic seep sediments. Comparisons are summarized for the depth intervals of 0 cmbsf (n=137 negative, n=9 biogenic, and n=8 thermogenic), 20-50 cmbsf (n=80 negative, n=9 biogenic, n=9 thermogenic), and >50 cmbsf (n=254 negative, n=29 biogenic, n=14 thermogenic). The IndicSpecies Stat value comparing nonseep with thermogenic or biogenic seeps for each indicator OTU is included, where a higher value indicates a stronger association.

Caldatribacteriota OTUs 1, 3029, and 21130, which belong to the JS1 lineage (genus-level), exhibited significantly different (p<0.05) relative sequence abundances in sediments with thermogenic hydrocarbons compared to non-seep sediments. OTU 3029 was significantly different across all depth categories, whereas OTU 1 was significantly different in 0 and >50 cmbsf sediments, and OTU 21130 was significantly different in 20-50 cmbsf and >50 cmbsf sediments (Table S1). In all sediments regardless of hydrocarbon ranking, a general trend of increasing relative sequence abundance of Caldatribacteriota OTUs with sediment depth was observed, with OTU 3029 showing the greatest increase with depth in sediments containing thermogenic hydrocarbons (Figure 3).

Campilobacterota OTU 8, affiliated with the genus Sulfurovum, was also detected in significantly greater (p<0.05) relative sequence abundance in sediments with thermogenic hydrocarbons compared to non-seep sediments at 0 cmsbf and 20-50 cmbsf (Table S1). OTU 8 was detected in greatest relative sequence abundance at 20-50 cmbsf in thermogenic hydrocarbon seep sediments, whereas in non-seep sediments, its sequence abundance was consistently low across all depths (Figure 3).

Acidobacteriota OTU 21, belonging to the Aminicenantales lineage, was detected in significantly different (p<0.05) relative sequence abundances in sediments with thermogenic hydrocarbons compared to non-seep sediments, most notably at 20-50 cmbsf (Figure 3).

Two of the five indicator OTUs for thermogenic hydrocarbons (Campilobacterota OTU 8 and Caldatribacteriota OTU 3029) were also detected at elevated levels in sediments with evidence of biogenic hydrocarbon seepage. In addition, all three Campilobacterota indicator OTUs (8, 9 and 47) were detected in higher relative sequence abundance in biogenic hydrocarbon seep sediments across all depths (Figure 3). These OTUs were detected in greatest relative sequence abundances at 20-50 cmbsf in both biogenic and thermogenic seep sites.

For these OTUs to have utility as hydrocarbon seepage indicators, their differential occurrence in thermogenic and biogenic sites compared to non-seep sites should be easily distinguishable. The seven indicator OTUs described above show the largest differences in this regard (on average >5%). Therefore, these OTUs were assessed in more detail to gain a better understanding of sites that were inconclusive based on geochemistry. To do this, OTUs 1 and 21 were used as exclusive indicators of thermogenic hydrocarbons and the remaining five OTUs, 8, 9, 47, 3029, and 21130, as shared indicator OTUs for thermogenic or biogenic hydrocarbon seepage.

### 3.5 Bacterial community analysis reveals sites with inconclusive geochemistry that warrant further investigation

Of the five sites with inconclusive geochemical evidence for thermogenic or biogenic hydrocarbon seepage (Table 1), three sites (15-6, 16-4, and 16-21) had bacterial communities that are significantly different from non-seep sites than sites with thermogenic or biogenic hydrocarbons, based on ANOSIM of entire bacterial communities (Table S7), when depth interval categories were considered separately. The 0 cmbsf bacterial community from inconclusive site 16-21 exhibited similarity to those in biogenic sediments, while its 20-50 cmbsf community was more similar to thermogenic sediments. For >50 cmbsf sediment, inconclusive site 15-6 was significantly different from non-seep sediments than both thermogenic and biogenic hydrocarbon seeps. Also at this depth, inconclusive site 16-4 was not significantly different from biogenic hydrocarbon seep sediments. Despite partial geochemical evidence for the presence of thermogenic hydrocarbons, sediments from inconclusive sites 15-9 and 16-32 do not have a bacterial signature that is suggestive of the presence of thermogenic or biogenic hydrocarbons.

The seven OTUs described above were used to assess inconclusive sites 15-6, 16-4, and 16-21 in more detail. Sediments from site 16-21, where geochemical analysis suggested hydrocarbons of both biogenic and thermogenic origin, exhibit increased relative sequence abundances of all seven indicator OTUs across the depth intervals. Six of the seven indicator OTUs are detected in higher relative sequence abundances at 20-50 cmbsf compared to non-seep sites (Figure 4). Inconclusive site 15-6 sediments have isotopic signatures suggestive of mixed biogenic-thermogenic origin (Table 1) and show greater relative sequence abundances of Caldatribacteriota OTUs 1, 3029 and 21130 at all depths, when compared to non-seep sediments. On the other hand, site 15-6 does not feature elevated abundances of Campilobacterota OTUs 8, 9 and 47 and Acidobacteriota OTU 21. Similarly, at site 16-4 indicator Campilobacterota OTUs 8, 9, and 47 were detected in very low relative abundances at all depths, where geochemical assessments pointed to only thermogenic hydrocarbon gas. However, this site features Caldatribacteriota OTUs 1 and 3029 in increased relative sequence abundance at >50 cmbsf, as observed in thermogenic seep sites.

**Figure 4.**
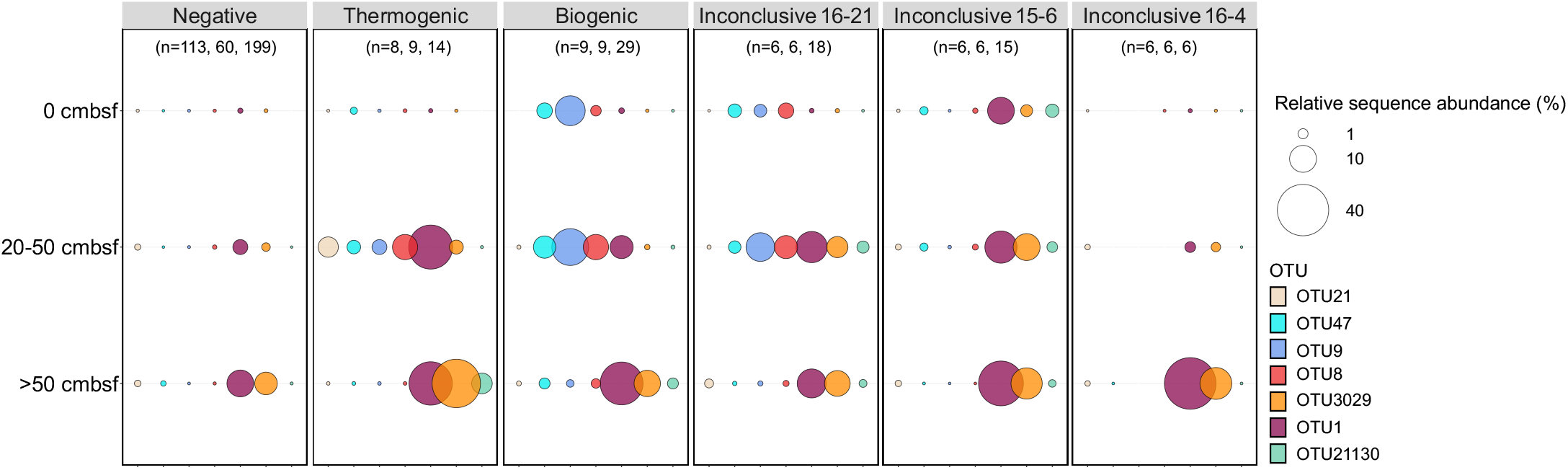
Comparison of average relative sequence abundances of indicator OTUs for thermogenic and biogenic hydrocarbon seepage in non-seep, thermogenic, biogenic and inconclusive sites at 0 cmbsf, 20-50 cmbsf, and >50 cmbsf. Number of replicates (n) are indicated for 0 cmbsf, 20-50 cmbsf, and >50 cmbsf depths, respectively. Thermogenic and biogenic hydrocarbon seep indicators Aminicenantales OTU 21, Campilobacterota OTUs 8, 9, 47 and Caldatribacteriota OTUs 1, 3029, and 21130 are represented by different colours and their relative sequence abundance is represented by the size of the circle.

## 4. Discussion

### 4.1 Uncultured bacteria associate with biogenic and thermogenic hydrocarbon geofluids

Sulfurovum and Sulfurimonas are commonly detected in the deep sea, including in and around cold-seeps, hydrothermal vents, and in oxic surface sediments (Inagaki et al. 2004; Bourbonnais et al. 2012; Deja-Sikora et al. 2019). As putative sulfur- and sulfide-oxidizing bacteria (SOB), they can consume reduced sulfur compounds such as the H_2_S generated by sulfate-reducing bacteria (SRB). In 20-50 cmbsf seep sediments where OTUs of these putative SOB were detected in highest relative sequence abundance, especially in biogenic seep sediments, it is likely that Sulfurovum and Sulfurimonas are able to combine aerobic respiration or nitrate reduction with the oxidation of sulfide (Han & Perner, 2015). Sulfide is generated deeper in the sediments, where elevated levels of sulfate reduction are fuelled by the presence of hydrocarbons, including methane. Syntrophic consortia in these settings consist of archaea that oxidize methane and other hydrocarbons together with sulfate-reducing bacteria (SRB), such as members of the SEEP-SRB1 lineage. While archaea were not included in the present study, several SEEP-SRB1 OTUs were associated with biogenic seeps (Table S2). Sulfurovum have been found in methane seep sediments along with many of the other taxa identified in this study (Trembath-Reichert et al. 2016). In particular, SEEP-SRB1 and Caldatribacteriota OTUs are significantly associated with biogenic methane in Scotian Slope sediments.

Members of the phylum Caldatribacteriota (formerly named Atribacteria/JS1) are commonly reported as part of bacterial communities in subsurface environments, particularly in anoxic organic-rich marine sediments (Orcutt et al. 2011) and in methane hydrates, hydrocarbon seeps, and petroleum reservoirs (Dong et al. 2020; Webster et al. 2004; Inagaki et al. 2006; Pham et al. 2009; Orcutt et al. 2011; Kobayashi et al. 2012; Parks et al. 2014). It is generally thought that Caldatribacteriota are anaerobes that generate fermentation products such as acetate, ethanol and carbon dioxide that may in turn support other bacteria and methanogens (Carr et al. 2015; Lee et al. 2018). Recent metagenomic evidence suggests that some Caldatribacteriota possess genes involved in anaerobic hydrocarbon degradation (Liu et al. 2019; Sierra-Garcia et al. 2020), however the genomes analysed were derived from much deeper oil reservoir samples, and it is unclear if this is an important part of the ecophysiology of this largely uncultured group of bacteria (Katayama et al. 2020) in shallow seabed cold seeps.

Predicted physiological features of Caldatribacteriota do not offer an immediate explanation for their observation in near-surface hydrocarbon seep sediments and their co-occurrence with putative aerobic SOB like Sulfurovum and Sulfurimonas. Given Caldatribacteriota genotypes are suggestive of metabolism in the deeper anoxic layers of the marine subsurface where they are ubiquitous, it has been suggested that a combination of environmental selection and dispersal via geofluids contributes to their prevalence in shallower sediment layers at hydrocarbon seeps, as shown here (i.e., in 0 and 20-50 cmbsf depth categories) and elsewhere (Chakraborty et al. 2020). Upward transport of fluids has been invoked to explain observations of Caldatribacteriota at different seabed mud volcano sites (Hoshino et al. 2017; Ruff et al. 2019) as well as at gas seeps in the Gulf of Mexico (Chakraborty et al. 2020). Similar observations were made at the Southwest Indian Ridge spreading centre where a predominant subsurface Caldatribacteriota lineage was only detected in surface sediments associated with potential upward porewater flux, but not in background sediments away from the spreading ridge (Varliero et al. 2019).

An influence of advective geofluid transport can also explain distributions of Caldatribacteriota at different depth intervals in hydrocarbon-positive Scotian Slope sediments. Caldatribacteriota OTUs found throughout the deeper sediment layers were significantly enriched in shallow sediments (20-50 cmbsf) at seep sites, which may be due to transport during hydrocarbon migration. This enrichment pattern was not observed for Sulfurovum and Sulfurimonas OTUs. Following upward transport of Caldatribacteriota, their higher sequence abundance compared to other bacteria in shallow sediments (Figure 3) may additionally be due to a competitive advantage from being adapted to the stronger selective pressures of the deep subsurface. Caldatribacteriota have unique attributes that may provide advantages in the energy-limited deep subsurface (Bird et al. 2019) as well as a variety of other environmental stresses (Glass et al. 2021), including hypothesized behaviour of carbohydrate storage as so-called ‘selfish bacteria’ (Orsi 2018).

Differences in the potential ecological roles of Caldatribacteriota OTUs and Campilobacterota OTUs in thermogenic hydrocarbon seep sediments of the Scotian Slope are highlighted in geochemistry inconclusive sediments (Figure 4). Whereas the depth profiles of these seven indicator OTUs at site 16-21 could suggest a mixed biogenic-thermogenic hydrocarbon signature similar to some of the strong hydrocarbon seep sites, the absence of Campilobacterota indicator OTUs in site 16-4, where there is no isotopic evidence for biogenic gas, suggests that Sulfurovum and Sulfurimonas OTUs 8, 9, and 47 may be primarily associated with biogenic gas.

### 4.2 DNA based sediment surveys offer increased capacity to locate thermogenic seep sites

Geochemical measurements of seabed sediment core samples are the gold standard for detecting hydrocarbon seepage. While these tried and tested methods work well, it is not uncommon to encounter cases with inconclusive results, as highlighted by this study. While testing with this many geochemical parameters is advantageous in terms of generating multiple layers of evidence, the complexity of these analyses requires experienced interpretation for identifying thermogenic hydrocarbons and evaluating whether seabed thermogenic hydrocarbon seepage is occurring.

It should also be considered that migration pathways of hydrocarbons through marine sediments are not necessarily vertical and can occur at variable fluxes as macroseepage or microseepage (Abrams 2017). Piston-coring collects a vertical plug of marine sediment at a given location and may thus capture thermogenic hydrocarbon content in varying concentrations in some but not all sections of the core. In this study, bacterial community profiles and indicator OTUs in sediments with inconclusive geochemistry were observed in a range of depths. At inconclusive sites 16-21 and 15-6, thermogenic indicator OTUs were observed through the full depth profile (Figure 4) whereas at inconclusive site 16-4 thermogenic indicator OTUs were only observed in deeper sediments (>50 cmbsf). These observations combine to suggest that thermogenic hydrocarbon seep detection via sediment coring would benefit from replicate sampling and by systematically sampling along transects that move away from proposed seep sites.

DNA-based bacterial diversity analysis at thermogenic hydrocarbon seeps highlights a unique community profile characterized by a predominance of Caldatribacteriota, specifically members of its JS1 lineage, and of Campilobacterota, specifically the genus Sulfurovum. Where seepage of thermogenic hydrocarbons is occurring, Aminicenantales are also predominant (Figure 3). These same three bacterial groups were highlighted in a study of thermogenic seepage in the Gulf of Mexico (Chakraborty et al. 2020). Absence of strong signals for these lineages in non-seep sites (Figure 4) provides a basis for DNA sequencing to offer a layer of evidence additional to geochemistry, especially for evaluating sites where geochemical analyses are inconclusive. Of the five inconclusive sites in this study, two are predicted not to contain thermogenic hydrocarbons based on bacterial DNA analysis, whereas the other three are determined to be sites of interest (i.e., potentially containing thermogenic hydrocarbons) warranting further investigation.

The significance of bacterial indicator OTUs as being diagnostic for the presence of thermogenic hydrocarbons is most pronounced in shallow sediments just below the surface and within the top 50 cmbsf. This can be particularly advantageous for predicting the presence of thermogenic hydrocarbon seepage in instances where short cores are obtained, including gravity cores as in some of the sites in this study. In this depth range, differences in indicator OTUs and in overall bacterial community composition between thermogenic and biogenic seeps were most prominent, potentially allowing DNA sequencing to specifically point to deeply sourced hydrocarbon seepage of thermogenic origin.

## Acknowledgements

Special thanks to the operators and crew of the CCGS Hudson and Natural Resources Canada for sediment core collection and processing. Ship funding was provided by Nova Scotia Department of Natural Resources and Renewables, the Nova Scotia Offshore Energy Research Association, and Natural Resources Canada. The work was supported by research funding from Genome Canada’s Genomics Applications Partnership Program facilitated by Genome Atlantic and Genome Alberta (to CRJH and AM), the Canada Foundation for Innovation grant (CFI-JELF 33752 to CRJH) and a Campus Alberta Innovates Program chair (CRJH). The authors also wish to thank Carey Ryan and Rhonda Clark for research and logistics support.

## CRediT author statement

Carmen Li – Conceptualization; Methodology; Investigation; Validation; Formal analysis; Data curation; Writing-Original Draft; Visualization

Oyeboade Adebayo – Investigation

Deidra K. Ferguson – Investigation

Scott Wang – Investigation

Jayne E. Rattray – Writing-Review & Editing

Martin Fowler – Formal analysis; Writing-Review & Editing

Jamie Webb – Formal analysis

Calvin Campbell – Resources; Supervision; Project administration

Natasha Morrison – Resources; Project administration

Adam MacDonald – Resources; Supervision; Project administration

Casey R.J. Hubert – Conceptualization; Writing-Original Draft; Supervision; Funding acquisition

