## Supplementary Material for "Bacterial anomalies in seabed sediments associated with deep water hydrocarbon seepage"

**Supplementary Table 1.** OTUs associated with thermogenic hydrocarbon seep samples as determined by IndicSpecies analysis (p = 0.05, association function =r.g.). The specific depth category (0 cmbsf, 20-50 cmbsf, or >50 cmbsf) to which an OTU was associated is listed except where an OTU had no association (N/A) with a specific depth. OTUs listed were detected in at least 5% relative sequence abundance in at least one bacterial amplicon library (n = 621). OTUs in bold text had a >5% difference in average sequence abundance between non-seep and thermogenic hydrocarbon seep samples in at least one depth category. Asterisks (*) denote that the OTU was in a significantly different relative sequence abundance in that depth category in thermogenic hydrocarbon seep samples compared to non-seep samples, as determined by two-sample T-tests assuming unequal variance. OTUs that met both criteria were included in Figure 3.

| Phylum | OTU | Taxonomy | Associated depth (cmbsf) | Indic Species Stat | Indic Species P-value | Difference in average relative sequence abundance between non-seep and thermogenic seep samples (%) | | |
|---|---|---|---|---|---|---|---|---|
| | | | | | | 0 cmbsf | 20-50 cmbsf | >50 cmbsf |
| **Caldatribacteriota** | **3029** | **JS1** | 20-50 and >50 | 0.848 | 0.0067 | -0.08* | **26.66*** | **18.94*** |
| Campilobacterota | 47 | Sulfurovum | 20-50 | 0.669 | 0.0181 | 0.38 | 2.10* | -0.11 |
| Gammaproteobacteria | 126 | B2M28 | 0 | 0.924 | 0.0053 | 0.88* | 0.16* | 0.02 |
| Desulfobacterota | 54 | Desulfosarcinaceae | 20-50 | 0.734 | 0.0326 | -0.05* | 1.15* | -0.01 |
| **Campilobacterota** | **8** | **Sulfurovum** | 20-50 | 0.685 | 0.0104 | 0.03* | **8.64*** | 0.02 |
| Campilobacterota | 9 | Sulfurimonas | 20-50 | 0.761 | 0.0127 | 0.01 | 2.61* | 0.02* |
| **Caldatribacteriota** | **21130** | **JS1** | >50 | 0.753 | 0.0186 | -0.00 | 0.01* | **5.57*** |
| Gammaproteobacteria | 417 | Thiohalophilus | 0 and 20-50 | 0.643 | 0.0171 | 0.06 | 0.23* | 0.01 |
| Desulfobacterota | 5829 | SEEP-SRB1 | >50 | 0.342 | 0.0195 | 0.00 | 0.00 | 0.02 |
| Chloroflexi | 38 | SCGC-AB-539-J10 | >50 | 0.531 | 0.0435 | -0.01 | 1.33* | 2.80* |
| **Acidobacteriota** | **21** | **Aminicenantales** | 20-50 | 0.621 | 0.0142 | -0.01 | **5.15*** | -0.28* |
| **Caldatribacteriota** | **1** | **JS1** | >50 | 0.836 | 0.0184 | -0.06* | 1.47 | **25.03*** |
| Desulfobacterota | 104 | SEEP-SRB1 | >50 | 0.262 | 0.0390 | -0.00 | -0.00 | 0.52 |
| Alphaproteobacteria | 100 | Rhodobacteraceae | 20-50 | 0.748 | 0.0269 | -0.04* | 0.66* | -0.00 |
| Cloacimonadota | 307 | MSBL2 | >50 | 0.526 | 0.0093 | 0.00 | 0.00 | 1.55* |
| Actinobacteriota | 135 | FS118-23B-02 | >50 | 0.387 | 0.0179 | 0.00 | 0.00 | 0.05 |
| Bacteroidota | 19 | Bacteroidales | N/A | 0.139 | 0.0093 | -0.02* | -0.05* | 2.09 |
| Gammaproteobacteria | 820 | Enhydrobacter | >50 | 0.338 | 0.0180 | -0.00 | 0.01 | 1.19 |

**Supplementary Table 2.** OTUs associated with biogenic hydrocarbon seep samples as determined by IndicSpecies analysis (p = 0.05, association function =r.g.). The specific depth category (0 cmbsf, 20-50 cmbsf, or >50 cmbsf) to which an OTU was associated is listed except where an OTU had no association (N/A) with a specific depth. OTUs listed were detected in at least 5% relative sequence abundance in at least one bacterial amplicon library (n = 637). OTUs in bold text had a >5% difference in average sequence abundance between non-seep and biogenic hydrocarbon seep samples in at least one depth category. Asterisks (*) denote that the OTU was in a significantly different relative sequence abundance in that depth category in biogenic hydrocarbon seep samples compared to non-seep samples, as determined by two-sample T-tests assuming unequal variance. OTUs that met both criteria were included in Figure 3.

| Phylum | OTU | Taxonomy | Associated depth (cmbsf) | Indic Species Stat | Indic Species P-value | Difference in average relative sequence abundance between non-seep and thermogenic seep samples (%) | | |
|---|---|---|---|---|---|---|---|---|
| | | | | | | 0 cmbsf | 20-50 cmbsf | >50 cmbsf |
| **Caldatribacteriota** | **3029** | **JS1** | >50 | 0.768 | 0.0001 | 0.047 | 4.861* | **18.20*** |
| **Campilobacterota** | **47** | **Sulfurovum** | 20-50 | 0.758 | 0.0001 | 2.893* | **6.609*** | 1.109* |
| Gammaproteobacteria | 126 | B2M28 | 0 | 0.822 | 0.0001 | 2.127* | 0.840* | 0.030* |
| Desulfobacterota | 54 | Desulfosarcinaceae | 20-50 | 0.653 | 0.0001 | 0.060 | 1.418* | 0.663* |
| **Campilobacterota** | **8** | **Sulfurovum** | 20-50 | 0.924 | 0.0001 | 1.124* | **8.711*** | 0.842* |
| **Campilobacterota** | **9** | **Sulfurimonas** | 0 and 20-50 | 0.785 | 0.0001 | **12.613*** | **19.730*** | 0.458* |
| Caldatribacteriota | 21130 | JS1 | >50 | 0.635 | 0.0001 | 0.004 | 0.039* | 1.235* |
| Gammaproteobacteria | 417 | Thiohalophilus | 0 | 0.795 | 0.0001 | 2.730* | 1.009* | 0.044* |
| Desulfobacterota | 5829 | SEEP-SRB1 | >50 | 0.553 | 0.0001 | 0.006* | 0.626* | 2.504* |
| Desulfobacterota | 104 | SEEP-SRB1 | >50 | 0.522 | 0.0002 | -0.002* | -0.002 | 1.039* |
| Alphaproteobacteria | 100 | Rhodobacteraceae | 0, 20-50, and >50 | 0.559 | 0.0001 | 0.613* | 1.242* | 0.879* |
| Cloacimonadota | 307 | MSBL2 | >50 | 0.585 | 0.0001 | 0 | 0.001 | 0.041* |
| Actinobacteriota | 135 | FS118-23B-02 | >50 | 0.475 | 0.0006 | 0 | 0 | 0.011* |
| Bacteroidota | 19 | Bacteroidales | >50 | 0.715 | 0.0001 | -0.013* | 0.0180 | 0.844* |
| Bacteroidota | 148 | Bacteroidetes BD2-2 | 20-50 | 0.599 | 0.0001 | 0.311* | 1.144* | 0.392* |
| Desulfobacterota | 26 | SEEP-SRB1 | >50 | 0.550 | 0.0001 | 0.001 | 0.126 | 4.838 |
| Desulfobacterota | 201 | SEEP-SRB1 | 20-50 | 0.392 | 0.0016 | 0.004 | 0.903 | 0.322 |
| Acidobacteriota | 103 | Subgroup 21 | 0 and >50 | 0.191 | 0.0376 | 0.273* | 0.054* | 0.388 |
| Spirochaetota | 53 | Spirochaetaceae | >50 | 0.307 | 0.0156 | -0.027* | -0.089* | 0.604 |
| Gammaproteobacteria | 820 | Enhydrobacter | N/A | 0.098 | 0.0483 | -0.000 | 0 | 0.005 |

**Supplementary Table 3.** OTUs associated with hydrocarbon seep (biogenic and thermogenic hydrocarbon seeps) samples as determined by IndicSpecies analysis (p = 0.05, association function =r.g.). The specific depth category (0 cmbsf, 20-50 cmbsf, or >50 cmbsf) to which an OTU was associated is listed except where an OTU had no association (N/A) with a specific depth. OTUs listed were detected in at least 5% relative sequence abundance in at least one bacterial amplicon library (n = 668). OTUs in bold text had a >5% difference in average sequence abundance between non-seep and hydrocarbon seep samples in at least one depth category. Asterisks (*) denote that the OTU was in a significantly different relative sequence abundance in that depth category in hydrocarbon-seep samples compared to non-seep samples, as determined by two-sample T-tests assuming unequal variance.

| Phylum | OTU | Taxonomy | Associated depth (cmbsf) | Indic Species Stat | Indic Species P-value | Difference in average relative sequence abundance between non-seep and thermogenic seep samples (%) | | |
|---|---|---|---|---|---|---|---|---|
| | | | | | | 0 cmbsf | 20-50 cmbsf | >50 cmbsf |
| **Caldatribacteriota** | **3029** | JS1 | N/A | 0.481 | 0.0001 | -0.01 | **15.41*** | **17.61*** |
| Campilobacterota | 47 | Sulfurovum | 20-50 | 0.629 | 0.0001 | 1.71* | 4.35* | 0.69* |
| Gammaproteobacteria | 126 | B2M28 | 0 | 0.735 | 0.0001 | 1.54 | 0.50 | 0.03 |
| Desulfobacterota | 54 | Desulfosarcinaceae | 20-50 | 0.669 | 0.0001 | -0.01 | 1.26* | 0.55 |
| **Campilobacterota** | **8** | Sulfurovum | 20-50 | 0.767 | 0.0001 | 0.61 | **8.67*** | 0.57 |
| **Campilobacterota** | **9** | Sulfurimonas | N/A | 0.338 | 0.0001 | **6.68*** | **11.17*** | 0.31 |
| Caldatribacteriota | 21130 | JS1 | >50 | 0.572 | 0.0001 | 0.00 | 0.02 | 2.65* |
| Gammaproteobacteria | 417 | Thiohalophilus | 0 | 0.549 | 0.0001 | 1.47* | 0.62 | 0.03* |
| Desulfobacterota | 5829 | SEEP-SRB1 | >50 | 0.456 | 0.0001 | 0.00 | 0.31 | 1.70* |
| Desulfobacterota | 26 | SEEP-SRB1 | >50 | 0.448 | 0.0001 | 0.00 | 0.06 | 3.26* |
| Chloroflexi | 38 | SCGC-AB-539-J10 | N/A | 0.180 | 0.0004 | 0.00 | 0.73 | 0.68 |
| Acidobacteriota | 21 | Aminicenantales | 20-50 | 0.415 | 0.0001 | -0.01 | 2.47 | -0.22 |
| Spirochaetota | 53 | Spirochaetaceae | >50 | 0.247 | 0.0114 | -0.03 | -0.06 | 0.40 |
| **Caldatribacteriota** | **1** | JS1 | >50 | 0.645 | 0.0001 | -0.02 | 0.56 | **11.17*** |
| Bacteroidota | 148 | Bacteroidetes BD2-2 | 20-50 | 0.373 | 0.0004 | 0.00 | 0.65 | 0.26 |
| Gammaproteobacteria | 37 | Delftia | >50 | 0.405 | 0.0001 | 0.00 | 0.00 | 0.05 |
| Alphaproteobacteria | 11 | Allorhizobium-Neorhizobium-Pararhizobium-Rhizobium | >50 | 0.334 | 0.0006 | -0.01 | 0.00 | 0.08 |
| Firmicutes | 62 | Anaerobacillus | >50 | 0.326 | 0.0006 | -0.01 | 0.00 | 0.05 |
| Desulfobacterota | 104 | SEEP-SRB1 | >50 | 0.434 | 0.0001 | 0.00 | 0.00 | 0.87 |
| Alphaproteobacteria | 100 | Rhodobacteraceae | N/A | 0.408 | 0.0001 | 0.30 | 0.94 | 0.59 |
| Cloacimonadota | 307 | MSBL2 | >50 | 0.308 | 0.0007 | 0.00 | 0.00 | 0.53 |
| Actinobacteriota | 135 | FS118-23B-02 | >50 | 0.307 | 0.0003 | 0.00 | 0.00 | 0.02 |
| Desulfobacterota | 201 | SEEP-SRB1 | N/A | 0.171 | 0.0001 | 0.00 | 0.45 | 0.22 |
| Bacteroidota | 19 | Bacteroidales | >50 | 0.319 | 0.0001 | -0.02 | -0.02 | 1.26 |
| Alphaproteobacteria | 81 | Mesorhizobium | N/A | 0.137 | 0.0268 | 0.01 | 0.00 | 0.01 |
| Acidobacteriota | 103 | Subgroup 21 | N/A | 0.112 | 0.0001 | 0.14 | 0.07 | 0.25 |
| Gammaproteobacteria | 820 | Enhydrobacter | N/A | 0.111 | 0.0002 | 0.00 | 0.01 | 0.39 |
| Alphaproteobacteria | 89 | Sphingomonas | >50 | 0.245 | 0.0052 | 0.00 | 0.00 | 0.10 |

**Supplementary Table 4.** Two-sample T-test (assuming unequal variance) pairwise analyses of thermogenic hydrocarbon seep and non-seep sediment bacterial indicator OTUs at depths 0 cmbsf, ≤50 cmbsf, and >50 cmbsf. OTUs listed were detected in at least 5% relative sequence abundance in at least one amplicon library and were determined by IndicSpecies analysis to be significantly associated with thermogenic hydrocarbon seep sediment samples.

| Depth Category | OTUs with significant differences (p=0.05) | P-value | OTUs with no significant differences (p=0.05) | P-value |
|---|---|---|---|---|
| 0 cmbsf | JS1 OTU1 | 0.04969 | Sulfurimonas OTU9 | 0.07274 |
| | Sulfurovum OTU8 | 0.02161 | Aminicenantales OTU21 | 0.15476 |
| | Bacteroidales OTU19 | 0.00000 | SCGC-AB-539-J10 OTU38 | 0.25468 |
| | Desulfosarcinaceae OTU54 | 0.00004 | Sulfurovum OTU47 | 0.07804 |
| | Rhodobacteraceae OTU100 | 0.00900 | SEEP-SRB1 OTU104 | 0.12487 |
| | B2M28 OTU126 | 0.01578 | FS118-23B-02 OTU135 | N/A |
| | JS1 OTU3029 | 0.01873 | MSBL2 OTU307 | 0.35062 |
| | | | Thiohalophilus OTU417 | 0.09955 |
| | | | Enhydrobacter OTU820 | 0.08878 |
| | | | SEEP-SRB1 OTU5829 | N/A |
| | | | JS1 OTU21130 | 0.31909 |
| 20-50 cmbsf | Sulfurovum OTU8 | 0.02116 | JS1 OTU1 | 0.07998 |
| | Sulfurimonas OTU9 | 0.00700 | SEEP-SRB1 OTU104 | 0.12723 |
| | Bacteroidales OTU19 | 0.01658 | FS118-23B-02 OTU135 | N/A |
| | Aminicenantales OTU21 | 0.04394 | MSBL2 OTU307 | N/A |
| | SCGC-AB-539-J10 OTU38 | 0.00310 | Enhydrobacter OTU820 | 0.34659 |
| | Sulfurovum OTU47 | 0.00220 | SEEP-SRB1 OTU5829 | N/A |
| | Desulfosarcinaceae OTU54 | 0.00510 | | |
| | Rhodobacteraceae OTU100 | 0.00662 | | |
| | B2M28 OTU126 | 0.00914 | | |
| | Thiohalophilus OTU417 | 0.04169 | | |
| | JS1 OTU3029 | 0.00057 | | |
| | JS1 OTU21130 | 0.03063 | | |
| >50 cmbsf | JS1 OTU1 | 0.00000 | Sulfurovum OTU8 | 0.21855 |
| | Sulfurimonas OTU9 | 0.02148 | Bacteroidales OTU19 | 0.27126 |
| | Aminicenantales OTU21 | 0.00000 | Sulfurovum OTU47 | 0.08751 |
| | SCGC-AB-539-J10 OTU38 | 0.02263 | Desulfosarcinaceae OTU54 | 0.88849 |
| | MSBL2 OTU307 | 0.03153 | Rhodobacteraceae OTU100 | 0.49283 |
| | JS1 OTU3029 | 0.00000 | SEEP-SRB1 OTU104 | 0.31095 |
| | JS1 OTU21130 | 0.00065 | B2M28 OTU126 | 0.07464 |
| | | | FS118-23B-02 OTU135 | 0.12941 |
| | | | Thiohalophilus OTU417 | 0.27740 |
| | | | Enhydrobacter OTU820 | 0.18503 |
| | | | SEEP-SRB1 OTU5829 | 0.17981 |

**Supplementary Table 5.** Two-sample T-test (assuming unequal variance) pairwise analyses of biogenic hydrocarbon seep and non-seep sediment bacterial indicator OTUs at depths 0 cmbsf, ≤50 cmbsf, and >50 cmbsf. OTUs listed were detected in at least 5% relative sequence abundance in at least one amplicon library and were determined by IndicSpecies analysis to be significantly associated with biogenic hydrocarbon seep sediment samples.

| Depth Category | OTUs with significant differences (p=0.05) | P-value | OTUs with no significant differences (p=0.05) | P-value |
|---|---|---|---|---|
| 0 cmbsf | Sulfurimonas OTU9 | 0.023255 | JS1 OTU3029 | 0.41008 |
| | Sulfurovum OTU8 | 0.000397329 | SEEP-SRB1 OTU_26 | 0.345308299 |
| | Bacteroidales OTU19 | 0.018414 | JS1 OTU21130 | 0.28618298 |
| | Bacteroidetes_BD2-2 OTU_148 | 0.000506 | FS118-23B-02 OTU_135 | N/A |
| | Rhodobacteraceae OTU100 | 0.006538 | Desulfosarcinaceae OTU54 | 0.207924 |
| | B2M28 OTU126 | 0.000167 | MSBL2 OTU307 | N/A |
| | SEEP-SRB1 OTU104 | 0.007802 | Enhydrobacter OTU820 | 0.088783 |
| | Sulfurovum OTU47 | 0.000979 | SEEP-SRB1 OTU201 | 0.288187 |
| | SEEP-SRB1 OTU5829 | 0.028138 | | |
| | Subgroup_21 OTU_103 | 5.19E-05 | | |
| | Thiohalophilus OTU417 | 0.000461 | | |
| | Spirochaetaceae OTU_53 | 0.000748 | | |
| 20-50 cmbsf | Sulfurovum OTU8 | 1.93052E-05 | SEEP-SRB1 OTU104 | 0.340117 |
| | Sulfurimonas OTU9 | 3.93E-06 | FS118-23B-02 OTU135 | N/A |
| | SEEP-SRB1 OTU_26 | 0.011542141 | MSBL2 OTU307 | N/A |
| | SEEP-SRB1 OTU5829 | 0.010891 | Enhydrobacter OTU820 | N/A |
| | Subgroup_21 OTU_103 | 0.000184 | SEEP-SRB1 OTU201 | 0.079891 |
| | Sulfurovum OTU47 | 0.000207 | Bacteroidales OTU19 | 0.640576 |
| | Desulfosarcinaceae OTU54 | 0.00018 | | |
| | Rhodobacteraceae OTU100 | 0.003538 | | |
| | B2M28 OTU126 | 0.000352 | | |
| | Thiohalophilus OTU417 | 0.002369 | | |
| | JS1 OTU3029 | 0.0042 | | |
| | JS1 OTU21130 | 0.021950463 | | |
| | Spirochaetaceae OTU_53 | 6.41E-09 | | |
| | Bacteroidetes_BD2-2 OTU_148 | 3.62E-05 | | |
| >50 cmbsf | SEEP-SRB1 OTU_26 | 0.000637456 | Subgroup_21 OTU_103 | 0.277162 |
| | Sulfurimonas OTU9 | 8.9E-05 | Spirochaetaceae OTU_53 | 0.136172 |
| | Bacteroidales OTU19 | 3.77E-08 | SEEP-SRB1 OTU201 | 0.177803 |
| | Sulfurovum OTU8 | 5.92955E-05 | Enhydrobacter OTU820 | 0.352401 |
| | MSBL2 OTU307 | 0.000241 | | |
| | JS1 OTU3029 | 2.5E-08 | | |
| | JS1 OTU21130 | 5.11682E-05 | | |
| | Sulfurovum OTU47 | 0.000389 | | |
| | SEEP-SRB1 OTU104 | 0.001417 | | |
| | SEEP-SRB1 OTU5829 | 0.000264 | | |
| | Rhodobacteraceae OTU100 | 0.000915 | | |
| | Thiohalophilus OTU417 | 0.000297 | | |
| | B2M28 OTU126 | 0.000352 | | |
| | FS118-23B-02 OTU135 | 0.003985 | | |
| | Desulfosarcinaceae OTU54 | 0.001105 | | |
| | Bacteroidetes_BD2-2 OTU_148 | 0.001573 | | |

**Supplementary Table 6.** Two-sample T-test (assuming unequal variance) pairwise analyses of hydrocarbon seep (thermogenic hydrocarbon seep and biogenic hydrocarbon seep) and non-seep sediment bacterial indicator OTUs at depths 0 cmbsf, ≤50 cmbsf, and >50 cmbsf. OTUs listed were detected in at least 5% relative sequence abundance in at least one amplicon library and were determined by IndicSpecies analysis to be significantly associated with hydrocarbon seep sediment samples.

| Depth Category | OTUs with significant differences (p=0.05) | P-value | OTUs with no significant differences (p=0.05) | P-value |
|---|---|---|---|---|
| 0 cmbsf | Sulfurovum OTU8 | 0.00252 | Aminicenantales OTU21 | 0.050443 |
| | Sulfurimonas OTU9 | 0.02981 | SEEP-SRB1 OTU26 | 0.691317 |
| | Sulfurovum OTU47 | 0.00126 | Desulfosarcinaceae OTU54 | 0.861763 |
| | B2M28 OTU126 | 0.00002 | JS1 OTU3029 | 0.748461 |
| | Thiohalophilus OTU417 | 0.00266 | JS1 OTU21130 | 0.26472 |
| | SEEP-SRB1 OTU5829 | 0.03461 | JS1 OTU1 | 0.003794 |
| 20-50 cmbsf | Sulfurovum OTU8 | 0.00003 | Aminicenantales OTU21 | 0.061087 |
| | Sulfurimonas OTU9 | 0.00013 | JS1 OTU1 | 0.242373 |
| | SEEP-SRB1 OTU26 | 0.03188 | | |
| | Sulfurovum OTU47 | 0.00003 | | |
| | Desulfosarcinaceae OTU54 | 0.00000 | | |
| | B2M28 OTU126 | 0.00028 | | |
| | Thiohalophilus OTU417 | 0.00091 | | |
| | JS1 OTU3029 | 0.00050 | | |
| | SEEP-SRB1 OTU5829 | 0.01781 | | |
| | JS1 OTU21130 | 0.00928 | | |
| >50 cmbsf | JS1 OTU1 | 0.00001 | | |
| | Sulfurovum OTU8 | 0.00011 | | |
| | Sulfurimonas OTU9 | 0.00012 | | |
| | Aminicenantales OTU21 | 0.00217 | | |
| | B2M28 OTU126 | 0.00091 | | |
| | Sulfurovum OTU47 | 0.00230 | | |
| | Desulfosarcinaceae OTU54 | 0.00014 | | |
| | B2M28 OTU126 | 0.00014 | | |
| | Thiohalophilus OTU417 | 0.00027 | | |
| | JS1 OTU3029 | 0.00000 | | |
| | SEEP-SRB1 OTU5829 | 0.00040 | | |
| | JS1 OTU21130 | 0.00001 | | |

**Supplementary Table 7.** ANOSIM results of bacterial communities in thermogenic hydrocarbon seep, biogenic hydrocarbon seep, non-seep, and inconclusive sites at all depths and individual depths of 0 cmbsf, 20-50 cmbsf, and >50 cmbsf.

| Depth | Comparison | | ANOSIM Statistic | Significance value | Significantly different? (p = 0.05) | Conclusion |
|---|---|---|---|---|---|---|
| All | Non-seep | Thermogenic | 0.2538 | 0.0001 | Yes | Thermogenic hydrocarbon seep, biogenic hydrocarbon seep, and non-seep sediments are significantly different when all depths are considered.<br><br>Thermogenic hydrocarbon seep and biogenic hydrocarbon seep sediments are less different from one another than they each are from non-seep sediments.<br><br>All inconclusive sites are less different from non-seep sites than from thermogenic hydrocarbon seep sites.<br><br>All inconclusive sites are less different from non-seep sites than from biogenic hydrocarbon seep sites. |
| | | Biogenic | 0.2532 | 0.0001 | Yes | |
| | | 16-21 | 0.1307 | 0.0012 | Yes | |
| | | 15-9 | 0.0418 | 0.1636 | No | |
| | | 15-6 | 0.1035 | 0.0093 | Yes | |
| | | 16-4 | 0.1242 | 0.0055 | Yes | |
| | | 16-32 | 0.0702 | 0.0776 | No | |
| | Thermogenic | Biogenic | 0.1821 | 0.0004 | Yes | |
| | | 16-21 | 0.1830 | 0.0003 | Yes | |
| | | 15-9 | 0.2268 | 0.0001 | Yes | |
| | | 15-6 | 0.1568 | 0.0006 | Yes | |
| | | 16-4 | 0.2082 | 0.0031 | Yes | |
| | | 16-32 | 0.1767 | 0.0056 | Yes | |
| | Biogenic | 16-21 | 0.1941 | 0.0001 | Yes | |
| | | 15-9 | 0.3668 | 0.0001 | Yes | |
| | | 15-6 | 0.1525 | 0.0031 | Yes | |
| | | 16-4 | 0.4692 | 0.0001 | Yes | |
| | | 16-32 | 0.3597 | 0.0001 | Yes | |
| 0 cmbsf | Non-seep | Thermogenic | 0.0238 | 0.3889 | No | Thermogenic hydrocarbon seep and non-seep sediments are not significantly different at 0 cmbsf.<br><br>Biogenic hydrocarbon seep and non-seep sediments are significantly different at 0 cmbsf.<br><br>All inconclusive sites are less different from non-seep sites than from thermogenic hydrocarbon seep sites.<br><br>Though inconclusive site **16-21** is significantly |
| | | Biogenic | 0.6585 | 0.0001 | Yes | |
| | | 16-21 | 0.5850 | 0.0001 | Yes | |
| | | 15-9 | 0.0621 | 0.3128 | No | |
| | | 15-6 | 0.6650 | 0.0001 | Yes | |
| | | 16-4 | -0.0894 | 0.7228 | No | |
| | | 16-32 | -0.2602 | 0.9535 | No | |
| | Thermogenic | Biogenic | 0.9032 | 0.0001 | Yes | |
| | | 16-21 | 0.9934 | 0.0015 | Yes | |
| | | 15-9 | 0.9421 | 0.0007 | Yes | |
| | | 15-6 | 0.8120 | 0.0003 | Yes | |
| | | 16-4 | 0.4709 | 0.0025 | Yes | |
| | | 16-32 | 0.2849 | 0.0943 | No | |
| | Biogenic | 16-21 | 0.5192 | 0.0007 | Yes | |
| | | 15-9 | 0.9594 | 0.0004 | Yes | |
| | | 15-6 | 0.7146 | 0.0004 | Yes | |
| | | 16-4 | 0.9768 | 0.0004 | Yes | |
| | | 16-32 | 0.9088 | 0.0038 | Yes | |

| | | | | | | |
|---|---|---|---|---|---|---|
| | | | | | | different from both biogenic hydrocarbon seep and non-seep sites, it is less different from biogenic hydrocarbon seep sites than from non-seep sites at 0 cmbsf.<br><br>Inconclusive site 16-21 is less different from biogenic hydrocarbon seep sites than from thermogenic hydrocarbon seep sites at 0 cmbsf. |
| 20-50 cmbsf | Non-seep | Thermogenic | 0.8744 | 0.0001 | Yes | Thermogenic hydrocarbon seep, biogenic hydrocarbon seep, and non-seep sediments are significantly different at 20-50 cmbsf and ANOSIM values suggest the strongest difference at this depth compared to 0 cmbsf and >50 cmbsf.<br><br>Though inconclusive site **16-21** is significantly different from thermogenic hydrocarbon seep, biogenic hydrocarbon seep, and non-seep sites, it is less different from thermogenic hydrocarbon seep sites and biogenic hydrocarbon seep sites than from non-seep sites.<br><br>Inconclusive site 16-21 is less different from thermogenic hydrocarbon seep sites than from biogenic hydrocarbon seep sites at 20 cmbsf. |
| | | Biogenic | 0.9252 | 0.0001 | Yes | |
| | | 16-21 | 0.7884 | 0.0001 | Yes | |
| | | 15-9 | 0.4023 | 0.0082 | Yes | |
| | | 15-6 | 0.6158 | 0.0003 | Yes | |
| | | 16-4 | 0.3508 | 0.0187 | Yes | |
| | | 16-32 | 0.1109 | 0.2777 | No | |
| | Thermogenic | Biogenic | 0.9985 | 0.0002 | Yes | |
| | | 16-21 | 0.4924 | 0.0014 | Yes | |
| | | 15-9 | 1.0000 | 0.0003 | Yes | |
| | | 15-6 | 0.9412 | 0.0003 | Yes | |
| | | 16-4 | 0.8627 | 0.0004 | Yes | |
| | | 16-32 | 1.0000 | 0.0060 | Yes | |
| | Biogenic | 16-21 | 0.5556 | 0.0009 | Yes | |
| | | 15-9 | 1.0000 | 0.0001 | Yes | |
| | | 15-6 | 1.0000 | 0.0004 | Yes | |
| | | 16-4 | 0.9572 | 0.0001 | Yes | |
| | | 16-32 | 1.0000 | 0.0043 | Yes | |
| >50 cmbsf | Non-seep | Thermogenic | 0.4933 | 0.0001 | Yes | Thermogenic hydrocarbon seep, biogenic hydrocarbon seep, and non-seep sediments are significantly different at >50 cmbsf. |
| | | Biogenic | 0.5025 | 0.0001 | Yes | |
| | | 16-21 | 0.4175 | 0.0001 | Yes | |
| | | 15-9 | 0.0502 | 0.2377 | No | |
| | | 15-6 | 0.4009 | 0.0001 | Yes | |
| | | 16-4 | 0.2515 | 0.0225 | Yes | |

| | | | | | | |
|---|---|---|---|---|---|---|
| | | 16-32 | 0.2707 | 0.0020 | Yes | |
| | Thermogenic | Biogenic | 0.4328 | 0.0001 | Yes | Inconclusive site **15-6** is significantly different from thermogenic hydrocarbon seep, biogenic hydrocarbon seep, and non-seep sites but is less different from thermogenic hydrocarbon seep sites and from biogenic hydrocarbon seep sites than from non-seep sites at >50 cmbsf.<br><br>Inconclusive site 15-6 is less different from biogenic hydrocarbon seep sites than from thermogenic hydrocarbon seep sites.<br><br>Inconclusive site **16-4** is not significantly different from biogenic hydrocarbon seep sites at >50 cmbsf. |
| | | 16-21 | 0.8279 | 0.0001 | Yes | |
| | | 15-9 | 0.8165 | 0.0001 | Yes | |
| | | 15-6 | 0.3820 | 0.0001 | Yes | |
| | | 16-4 | 0.8863 | 0.0001 | Yes | |
| | | 16-32 | 0.6539 | 0.0001 | Yes | |
| | Biogenic | 16-21 | 0.6839 | 0.0001 | Yes | |
| | | 15-9 | 0.6592 | 0.0001 | Yes | |
| | | 15-6 | 0.2709 | 0.0003 | Yes | |
| | | 16-4 | 0.1036 | 0.1534 | No | |
| | | 16-32 | 0.6228 | 0.0001 | Yes | |

**Supplementary Table 8.** ANOSIM results of bacterial communities in hydrocarbon seep (thermogenic hydrocarbon seep and biogenic hydrocarbon seep), non-seep, and inconclusive sites at all depths and individual depths of 0 cmbsf, 20-50 cmbsf, and >50 cmbsf.

| Depth | Comparison | | ANOSIM Statistic | Significance value | Significantly different? (p = 0.05) | Conclusion |
|---|---|---|---|---|---|---|
| All | Non-seep | Positive | 0.2827 | 0.0001 | Yes | Hydrocarbon seep and non-seep sediments are significantly different when all depths are considered.<br><br>Inconclusive site **15-6** is significantly different from non-seep sites and not significantly different from hydrocarbon-seep sites.<br><br>Though inconclusive site **16-21** is significantly different from both hydrocarbon-seep and non-seep sites, it is the only 'inconclusive' location that is less different from hydrocarbon-seep sites than from non-seep sites. |
| | | 16-21 | 0.1670 | 0.0001 | Yes | |
| | | 15-9 | 0.0590 | 0.0701 | No | |
| | | 15-6 | 0.1377 | 0.0008 | Yes | |
| | | 16-4 | 0.1452 | 0.0025 | Yes | |
| | | 16-32 | 0.0948 | 0.0295 | Yes | |
| | | 16-5 | 0.1619 | 0.0011 | Yes | |
| | | 16-6 | 0.1199 | 0.0024 | Yes | |
| | | 16-13 | 0.0073 | 0.4119 | No | |
| | | 15-18 | 0.0595 | 0.0833 | No | |
| | Hydrocarbon seep | 16-21 | 0.1068 | 0.0104 | Yes | |
| | | 15-9 | 0.2892 | 0.0001 | Yes | |
| | | 15-6 | 0.0280 | 0.2453 | No | |
| | | 16-4 | 0.3164 | 0.0003 | Yes | |
| | | 16-32 | 0.2298 | 0.0001 | Yes | |
| | | 16-5 | 0.2382 | 0.0005 | Yes | |
| | | 16-6 | 0.3633 | 0.0001 | Yes | |
| | | 16-13 | 0.3894 | 0.0001 | Yes | |
| | | 15-18 | 0.4146 | 0.0001 | Yes | |
| 0 cmbsf | Non-seep | Positive | 0.3302 | 0.0005 | Yes | Hydrocarbon seep and non-seep sediments are significantly different at 0 cmbsf.<br><br>Though inconclusive sites **16-21** and **15-6** are significantly different from both hydrocarbon seep and non-seep sites, they are less different from hydrocarbon-seep sites than from non-seep sites. |
| | | 16-21 | 0.5373 | 0.0003 | Yes | |
| | | 15-9 | -0.0001 | 0.4702 | No | |
| | | 15-6 | 0.6352 | 0.0002 | Yes | |
| | | 16-4 | -0.0917 | 0.7404 | No | |
| | | 16-32 | -0.2189 | 0.9122 | No | |
| | | 16-5 | -0.2111 | 0.8991 | No | |
| | | 16-6 | -0.1680 | 0.9197 | No | |
| | | 16-13 | 0.0192 | 0.4028 | No | |
| | | 15-18 | -0.1667 | 0.9124 | No | |
| | Hydrocarbon seep | 16-21 | 0.1968 | 0.0284 | Yes | |
| | | 15-9 | 0.3220 | 0.0090 | Yes | |
| | | 15-6 | 0.5527 | 0.0003 | Yes | |
| | | 16-4 | 0.2640 | 0.0199 | Yes | |
| | | 16-32 | 0.0972 | 0.1506 | No | |
| | | 16-5 | 0.1076 | 0.1414 | No | |
| | | 16-6 | 0.1872 | 0.0458 | Yes | |
| | | 16-13 | 0.3129 | 0.0052 | Yes | |
| | | 15-18 | 0.1769 | 0.0502 | No | |
| 20-50 cmbsf | Non-seep | Positive | 0.9955 | 0.0001 | Yes | Hydrocarbon seep and non-seep sediments are significantly different at 20-50 cmbsf and ANOSIM values suggest the strongest difference at this depth |
| | | 16-21 | 0.9760 | 0.0001 | Yes | |
| | | 15-9 | 0.5845 | 0.0004 | Yes | |
| | | 15-6 | 0.8284 | 0.0001 | Yes | |
| | | 16-4 | 0.4777 | 0.0037 | Yes | |
| | | 16-32 | 0.2915 | 0.0950 | No | |

| | | | | | | |
|---|---|---|---|---|---|---|
| | | 16-5 | 0.9985 | 0.0001 | Yes | compared to 0 cmbsf and >50 cmbsf.<br><br>Though inconclusive sites **16-21** and **15-6** are significantly different from both hydrocarbon seep and non-seep sites, they are less different from hydrocarbon seep sites than from non-seep sites.<br><br>The highest ANOSIM values were from samples at this depth. |
| | | 16-6 | 0.7810 | 0.0001 | Yes | |
| | | 16-13 | 0.2672 | 0.476 | Yes | |
| | | 15-18 | 0.0931 | 0.2455 | No | |
| | Hydrocarbon seep | 16-21 | 0.2712 | 0.0250 | Yes | |
| | | 15-9 | 0.8952 | 0.0002 | Yes | |
| | | 15-6 | 0.7513 | 0.0001 | Yes | |
| | | 16-4 | 0.9645 | 0.0001 | Yes | |
| | | 16-32 | 0.8022 | 0.0005 | Yes | |
| | | 16-5 | 0.9993 | 0.0010 | Yes | |
| | | 16-6 | 0.9939 | 0.0001 | Yes | |
| | | 16-13 | 0.9688 | 0.0001 | Yes | |
| | | 15-18 | 0.9655 | 0.0001 | Yes | |
| >50 cmbsf | Non-seep | Positive | 0.6558 | 0.0001 | Yes | Hydrocarbon seep and non-seep sediments are significantly different at >50 cmbsf.<br><br>Inconclusive site **16-4** is significantly different from both hydrocarbon seep and non-seep sites but is less different from hydrocarbon seep sites than from non-seep sites. |
| | | 16-21 | 0.5632 | 0.0001 | Yes | |
| | | 15-9 | 0.1184 | 0.0420 | Yes | |
| | | 15-6 | 0.5376 | 0.0001 | Yes | |
| | | 16-4 | 0.3732 | 0.0007 | Yes | |
| | | 16-32 | 0.3837 | 0.0001 | Yes | |
| | | 16-5 | 0.3297 | 0.0001 | Yes | |
| | | 16-6 | 0.2420 | 0.0006 | Yes | |
| | | 16-13 | 0.0711 | 0.1697 | No | |
| | | 15-18 | 0.4653 | 0.0001 | Yes | |
| | Hydrocarbon seep | 16-21 | 0.6461 | 0.0001 | Yes | |
| | | 15-9 | 0.6872 | 0.0001 | Yes | |
| | | 15-6 | 0.1770 | 0.0085 | Yes | |
| | | 16-4 | -0.0592 | 0.7026 | No | |
| | | 16-32 | 0.6610 | 0.0001 | Yes | |
| | | 16-5 | 0.5129 | 0.0001 | Yes | |
| | | 16-6 | 0.7490 | 0.0001 | Yes | |
| | | 16-13 | 0.9048 | 0.0001 | Yes | |
| | | 15-18 | 0.8240 | 0.0001 | Yes | |
